# Wnt signaling couples G2 phase control with differentiation during hematopoiesis

**DOI:** 10.1101/2023.06.29.547151

**Authors:** Lauren M. Goins, Juliet R. Girard, Bama Charan Mondal, Sausan Buran, Chloe C. Su, Ruby Tang, Titash Biswas, Utpal Banerjee

## Abstract

During homeostasis, a critical balance is maintained between myeloid-like progenitors and their differentiated progeny, which function to mitigate stress and innate immune challenges. The molecular mechanisms that help achieve this balance are not fully understood. Using genetic dissection in *Drosophila*, we show that a Wnt6/EGFR-signaling network simultaneously controls progenitor growth, proliferation, and differentiation. Unlike G1-quiescence of stem cells, hematopoietic progenitors are blocked in the G2 phase by a β-catenin-independent Wnt6 pathway that restricts Cdc25 nuclear entry and promotes cell growth. Canonical β-catenin-dependent Wnt6 signaling is spatially confined to mature progenitors through localized activation of the tyrosine-kinases EGFR and Abl, which promote nuclear entry of β-catenin and facilitate exit from G2. This strategy combines transcription-dependent and - independent forms of both Wnt6 and EGFR pathways to create a direct link between cell-cycle control and differentiation. This unique combinatorial strategy employing conserved components may underlie homeostatic balance and stress response in mammalian hematopoiesis.

## Introduction

Precise spatiotemporal control of the cell cycle is a critical component of normal development across the evolutionary spectrum. Intrinsic control of cell cycle machinery follows a well-defined series of regulated activation and degradation of its core components^1–3^. However, within a developing tissue, the cell cycle is also tightly controlled by the spatial and temporal context of the cell. Individual cells divide and differentiate at their own unique rates, and it is expected that extracellular signals will coordinate the timing and frequency of entry into, and exit from, the mitotic cycle^4, 5^. For example, external signals perceived in G1 typically dictate whether a cell should self-renew, differentiate, or remain quiescent^6–9^. In the past, suggestions that cell cycle and differentiation are linked have largely been proposed in the context of the events restricted to G1^10–12^. Examples of regulation at the G2 stage are rarer and less well understood. In mammalian systems, a “G2 arrest” is generally associated with radiation-induced DNA damage, which is sensed by the ATM/ATR group of proteins^13, 14^. Under radiation stress, cells are held in G2 stasis until a repair mechanism is able to reverse the damage. During normal development, without induced DNA damage, a cell in G2 with 4N DNA content will not interpret differentiation signals before it passes the intervening step of mitosis. However, this does not imply that passage through G2 is a passive process, incapable of reacting to external signals. In fact, such extrinsic signals often inform the fates of the immediate daughter cells, such as during asymmetric cell division^15, 16^.

Here, we explore a molecular mechanism that links the progression of a cell from G2 into mitosis with the differentiation of its daughter cells during the following G1 phase of the cell cycle. In brief, we demonstrate that in an early, proliferative stage in development, cells self-renew and increase in number, whereas at a more mature stage, a regulated G2 phase block, enforced in the progenitor, plays a pivotal role in the decision of its daughter cells to choose differentiation over proliferation. The underlying mechanism involves multiple forms of Wnt signaling and multiple targets of tyrosine kinases in the context of E-Cadherin expression. A similar convergence of signals could control mammalian multipotent progenitors as they differentiate into cells of the myeloid lineage.

Several alternative modes of Wnt signaling have been described in the literature^17^. One such form, referred to as “WNT-STOP”, is most apparent in G2^18–21^. During this cell cycle phase, inhibition of GSK3 by the Wnt signal prevents a large and varied number of its target proteins from degradation. The resulting accumulation of protein mass is vital for the growth of the cell during G2 and a loss of this form of the Wnt signal therefore results in a smaller cell size. Although β-catenin is also stabilized by inhibition of GSK3, the cell-growth-related activity of Wnt is β-catenin independent. In what is termed “canonical Wnt signaling”, stabilized β-catenin, upon further modifications, enters the nucleus, and participates in transcriptional control of downstream targets^22^.

Two separate populations of β-catenin have been identified in the cell, the first is in a complex that includes E-Cadherin and α-catenin and is restricted to the cell surface and located at junctions between cells^23, 24^. The second population comprises a specifically phosphorylated form of β-catenin that is targeted to the nucleus to function in a transcription complex with TCF/LEF1^22^. Both sub-populations of β-catenin are stabilized against degradation by Wnt signaling^25^. Here we establish that interconnected β-catenin-independent and β-catenin-dependent Wnt pathways play a major role in regulating the spatial and temporal control of hematopoietic progenitors. This process directly links the cell cycle status of the progenitors with the cell fate choice of their progeny.

The lymph gland is the primary hematopoietic organ in *Drosophila*. Many similarities with mammalian myeloid hematopoiesis have made this tissue useful as an *in vivo* genetic model^26, 27^. Specified in the late embryo, the lymph gland develops during the larval instars and disintegrates at the pupal stage to release hemocytes that populate both the pupa and the adult^28, 29^. Functional zones of cells with distinguishing characteristics have been identified and characterized in the lymph gland^30^ (Figure 1A). The cells of the Posterior Signaling Center (PSC), along with similar cells along the dorsal vessel (heart), function as a niche to maintain progenitors in the medullary zone (MZ), preventing them from prematurely differentiating into hemocytes^31–35^.

**Figure 1:**
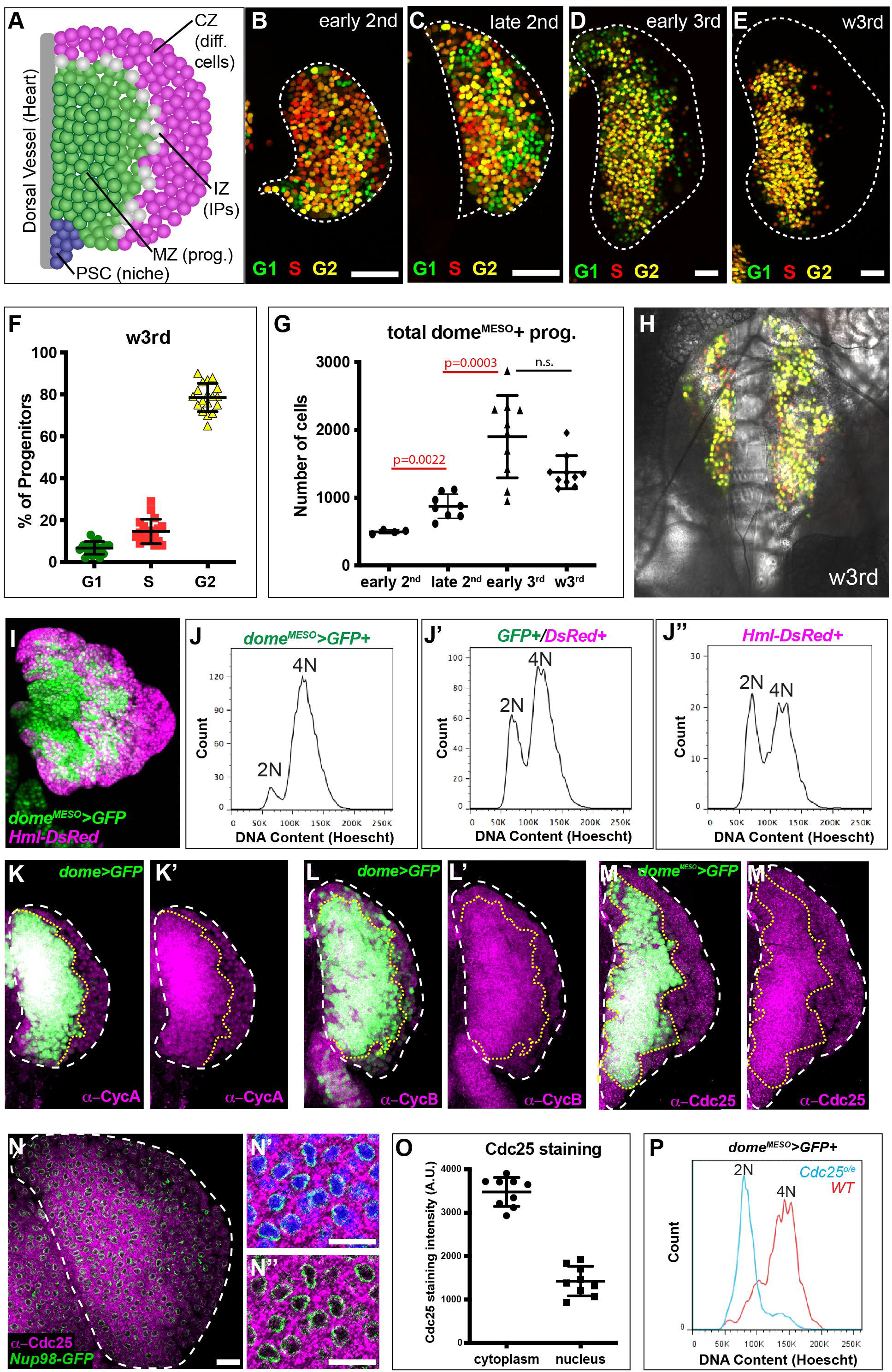
Hematopoietic progenitors are held in G2 phase of the cell cycle through sequestration and inactivation of Cdc25/String. **(A)** Schematic representation of functional zones of cells in the wandering third (w3rd) instar larval lymph gland (LG). The gray tube represents the dorsal vessel or heart, to which the LG lobes are attached. One lobe of the lymph gland is shown here and in the other panels. The Posterior Signaling Center (PSC; blue) consists of a population of cells with properties of a hematopoietic niche. The adjacent Medullary Zone (MZ) houses a heterogeneous group of hematopoietic progenitors (prog.; green). The Cortical Zone (CZ) contains differentiated hemocytes (diff. cells; magenta) and this zone continues to the distal periphery of the LG. Juxtaposed between the MZ and CZ, a small population of cells labeled Intermediate Progenitors (IPs; white) have transitional properties and belong to a zone named the Intermediate Zone (IZ). **(B-H)** A *GAL4/UAS* driven Fly-FUCCI system is used to monitor the cell cycle status of individual progenitors (genotype: *dome^MESO^-GAL4*; *UAS-FUCCI*). GFP marks cells in G1 (green). S phase (red) cells are marked by RFP. G2 (yellow) cells express both GFP and RFP. **(B-E)** Developmental stage-dependent FUCCI expression in the progenitor population. **(B)** early 2nd, **(C)** late 2nd, **(D)** early 3rd, and **(E)** w3rd instar larvae. Most *dome^MESO^*-expressing progenitors are in G2 by the 3rd instar. The outline of the entire LG lobe (white dashes) is based on DAPI staining omitted here for clarity. **(F)** Quantitative analysis of FUCCI expression in individual progenitors of w3rd instar larvae. Nearly 80% of the progenitors are in G2, with a small percentage in S (13%) and in G1 (8%). **(G)** Quantitative analysis of total progenitor number over developmental time. The number of progenitors significantly increases from early 2nd to late 2nd instar and from late 2nd to early 3rd, but there is no significant change in the overall progenitor number between early 3^rd^ and w3rd instar larvae. **(H)** Live imaging of a w3rd instar LG with progenitors expressing FUCCI. This panel shows a single slice from a z-stack (please see **Movie 1** for a view of the entire LG). This image illustrates the preponderance of G2 (yellow) cells at this stage. The transmitted light image (gray) is superimposed to show the rest of the LG cells that are not progenitors as well as other associated larval tissues. **(I)** A representative image of a LG with marked progenitors (*dome^MESO^>GFP*; green), differentiating cells (*Hml-DsRed*; magenta), and double positive intermediate progenitors (white due to the combination of green and magenta). **(J-J’’)** Flow cytometric DNA content analysis (Hoescht) of dissociated cells from lymph glands with the genotype in **(I).** Most progenitors **(J)** have 4N DNA content (G2 phase), whereas significant numbers of intermediate progenitors **(J’)** and differentiating cells **(J’’)** show a mixture of 2N (G1) and 4N (G2) DNA content suggesting that these populations are no longer blocked in G2. **(K-M’)** High levels of G2 cyclins, Cyclin A (magenta; **K, K’**) and Cyclin B (magenta; **L, L’**), as well as the Cdc25 phosphatase String (magenta; **M, M’**) are seen in progenitors (GFP, green; demarcated by yellow dotted line). GFP expression is controlled by *dome^MESO^-GAL4* **(M, M’)** or the related *domeless-GAL4* **(K-L’)**. **(N-N”)** *Nup98-GFP* (green) marks the nuclear envelope and DAPI (blue; **N’**) marks DNA. The lower **(N)** and higher **(N’, N”)** magnification views of the same LG lobe illustrate that Cdc25/String (magenta) is excluded from the nucleus and restricted to the cytoplasm. **(O)** Quantification of Cdc25/String staining data shown in **(N)**. **(P)** Overexpression of Cdc25 in the progenitors (*UAS-Cdc25*) causes a dramatic change in their cell cycle profile from predominantly in G2 (4N) in WT (red) to G1 (2N) in *UAS-Cdc25* (blue). See also **Movie 1** and **Figure S1**.

Recent genetic^36–39^ and transcriptomic studies^40, 41^ have identified considerable heterogeneity within the progenitor population. These subpopulations are spatially arranged so that the innermost ones (“core” progenitors), closest to the dorsal vessel, are the least mature, while the most mature ones (“distal” progenitors) are found at the periphery of the MZ near the site of differentiation. A transition zone, made up of intermediate progenitors^42^ (IPs) as well as other less well-characterized transitional cells^40, 41^, mark the “edge” where differentiation is first seen. Final maturation and lineage choice are largely restricted to the cortical zone (CZ), which extends to the periphery of the lymph gland and is composed of multiple mature cell types.

The proliferation profile of hematopoietic progenitors is quite distinct between early *vs* later larval stages. In the 1^st^ instar, the progenitors proliferate quite broadly, and this expansion phase continues until the mid 2^nd^ instar^32, 43^. At this point, two important and virtually simultaneous events become apparent. The first is the appearance of a small number of differentiating cells and the second is a dramatic reduction in proliferation of the progenitors. The proliferation block is enforced by two signals, a Hedgehog-dependent signal from the niche^33^ and a second, ADGF/Adenosine-based equilibrium signal from the early differentiating cells to the progenitors^43^. Together they suppress the proliferation of progenitors, whereas differentiation is confined to the distal edge of the MZ.

Differentiation is defined both by a reduction in the level of progenitor markers and by the appearance of a marker such as Hemolection (Hml), which identifies cells that have become competent to form all mature blood cell types and is expressed earlier than markers for terminal differentiation. The appearance of Hml correlates with the slowing down of progenitor proliferation such that by the 3^rd^ instar, where most of this study is focused, very few progenitors incorporate BrdU^32^. Additionally, differentiation is limited to the outer edge of the MZ, and processes that control proliferation and differentiation together maintain a balanced and spatially organized population of cells in each zone^30^. This orderly homeostatic control is disrupted upon immune challenge when progenitors rapidly proliferate and differentiate to protect the animal^44, 45^. In this study, we focus on mechanisms that are crucial for linking proliferative and cell-fate determinative events during normal hematopoiesis.

## Results

### Hematopoietic progenitors are held in the G2 phase of the cell cycle

We first sought to determine the cell cycle status of the slow-cycling hematopoietic progenitors. Recent studies have found that the hematopoietic progenitors in the lymph gland are largely in the G2 phase, although this varies by developmental time and the specific driver used to mark the cells (^46, 47^). Here, we expressed a fluorescent ubiquitin-based cell cycle indicator system (Fly-FUCCI^48^) in the progenitors using *dome^MESO^-GAL4* to identify and visualize cells in G1, S, and G2 phases of the cell cycle (Figure 1B-E). We find that a vast majority (∼80%) of the progenitors in the wandering 3^rd^ instar (w3^rd^ instar) lymph gland are held in G2 (Figure 1E-F), quite unlike classically defined “quiescent” cells in other developmental systems that are held in G1 or G0^8^. This “G2 block” or lengthening of the G2 phase is seen as early as 2^nd^ instar although it is most apparent in w3^rd^ (Figure 1B-F). Analysis of fixed tissue shows that progenitor number increases during the 2^nd^ instar but remains relatively constant throughout the 3^rd^ (Figure 1G). Mitotic cells are detected throughout the progenitor population in the 2^nd^ instar (Figure S1A,A’). These events are rarer in the 3^rd^ instar, and the ones detected are generally confined to the distal edge of the progenitor population near the site of differentiation (Figure S1B,B’). We devised a single-cell resolution, short time span, live imaging method, which, in combination with FUCCI, allows us to detect the cell cycle status of individual progenitors. Amongst other advantages, this method of imaging preserves the native 3D spatial structure of the lymph gland and allows its analysis at specific time points in development. As expected, most progenitors detected are in G2 irrespective of the developmental stage and this is particularly obvious in w3^rd^ instar larvae (Figure 1H; please see Movie 1; Figure S1C-F).

Additionally, in the late 2^nd^ and early 3^rd^ instars, G1 cells appear clustered along the distal edge of the MZ (Figure S1G). The immediate neighbors of these G1 cells no longer express the progenitor marker *dome^MESO^* (reported by FUCCI), suggesting that the G1 region is close to sites of differentiation. In principle, the band of G1 daughter cells in the early 3^rd^ could potentially choose between self-renewal and differentiation. However, since we no longer detect a significant increase in the progenitor population past this stage (Figure 1G), we believe that self-renewal at the edge is less prevalent than differentiation into a *Hemolectin (Hml)*-positive cell.

Further insight into the cell cycle is obtained in flow cytometric analysis of dissociated cells from multiply marked lymph glands (*dome^MESO^-GFP, Hml-DsRed*) (Figure 1I-J”; Figure S1H). In this analysis, progenitors (GFP-positive) are largely 4N in their DNA content (G2 phase), whereas the intermediate progenitors (expressing both GFP and RFP) and the differentiating cells (RFP-positive) consist of a mixture of 2N and 4N DNA profiles indicating that they are in multiple stages of the cell cycle. This agrees with the FUCCI results and further reinforces the possibility of an active mechanism that holds a majority of the progenitors in G2.

### Progenitors show high levels of cytoplasmic Cdc25 and Cyclins A and B

Both classic G2 cyclins, Cyclin A (Figure 1K-K’) and Cyclin B (Figure 1L-L’), are enriched in progenitors. In *Drosophila*, G2 arrest has been previously described in the context of development and is generally associated with an absence of the phosphatase Cdc25 (*Drosophila* String), either due to the degradation of the protein or a lack of transcription of the *cdc25* gene^49–51^. Cdc25 normally activates the Cdk1/Cyclin B complex by removing an inhibitory phosphate group on CDK1^52^. This event is a prerequisite for exit from G2 and a lack of Cdc25 would therefore result in a G2 block.

Surprisingly, we find that Cdc25 protein is expressed in abundance in the hematopoietic progenitors that nevertheless remain held in G2 (Figure 1M-M’). This seemingly contradictory result is resolved by a closer examination of the Cdc25 staining pattern, which reveals that virtually all of the Cdc25 protein in the progenitors is excluded from the nucleus and is instead held in the cytoplasm (Figure 1N-O), where Cdc25 will not be able to act on its nuclear targets^53^. These observations suggest that an active molecular mechanism sequesters and retains Cdc25/String in the cytosol, which renders Cdc25 inactive and thereby holds the cell in G2. It is possible to override such a mechanism by gross overexpression of Cdc25/String. Doing so decreases the fraction of progenitors held in G2 (4N) and increases the fraction of progenitors in G1 (2N) (Figure 1P). This result is further confirmed by FUCCI analysis (Figure S1I-K).

### Mechanism of Cdc25 sequestration

To initiate a dissection of the mechanisms that underlie lengthened G2, we first manipulated Myt1 and Wee1, which are core components of the cell cycle machinery that holds cells in G2. Previously validated RNAis which target transcripts encoding Myt1 and Wee1 kinases were used. It is widely established that these kinases sequentially phosphorylate and inactivate the Cyclin B/Cdk1 complex^52, 54^. We find that a kno^32^ck-down of these kinases within the progenitors causes a dramatic reduction in their population with a concomitant increase in differentiation (Figure S2A-C). The progenitors that remain in these genetic backgrounds are altered in their cell cycle status with a significantly smaller fraction in G2 (Figure S2D-H). Thus, holding these cells in G2 is important for maintaining a reserve pool of progenitors and a release from G2 strongly correlates with increased differentiation.

In mammalian systems, G2 arrest is associated with radiation-induced DNA damage and the protein cascades that control repair are well established^55^. Although the G2 arrest in the hematopoietic progenitors described here is seen under normal growth conditions with no exposure to radiation, we nevertheless investigated if any of the key components of DNA-damage-induced G2 arrest might be involved in the context of the lymph gland. We find that loss of function of the ATR kinase (a DNA damage sensor^56, 57^) or its substrate, the Chk1 kinase (that phosphorylates Cdc25^14, 58–60)^, as well as the Cdc25 binding protein 14-3-3ε ^61, 62^, results in phenotypes similar to the loss of Wee1 and Myt1. In each case, progenitors are depleted, differentiation is increased, and the progenitors that remain lose their strict restriction for G2 (Figures 2A-D, S2D-E, S2I-K). Additionally, in these knock-down backgrounds, Cdc25/String staining is greatly reduced in the progenitors that remain (Figure 2E-H’). Mechanistically, the data, interpreted in the context of published literature on mammalian systems^63^, is consistent with the model that ATR activates Chk1 which then phosphorylates Cdc25 and inactivates it due to its binding to 14-3-3ε ^52^, thus preventing Cdc25 from entering the nucleus^64^. This pathway that is typically utilized in radiation-stress sensing is co-opted during homeostatic hematopoietic development and helps hold progenitors in their undifferentiated state.

**Figure 2:**
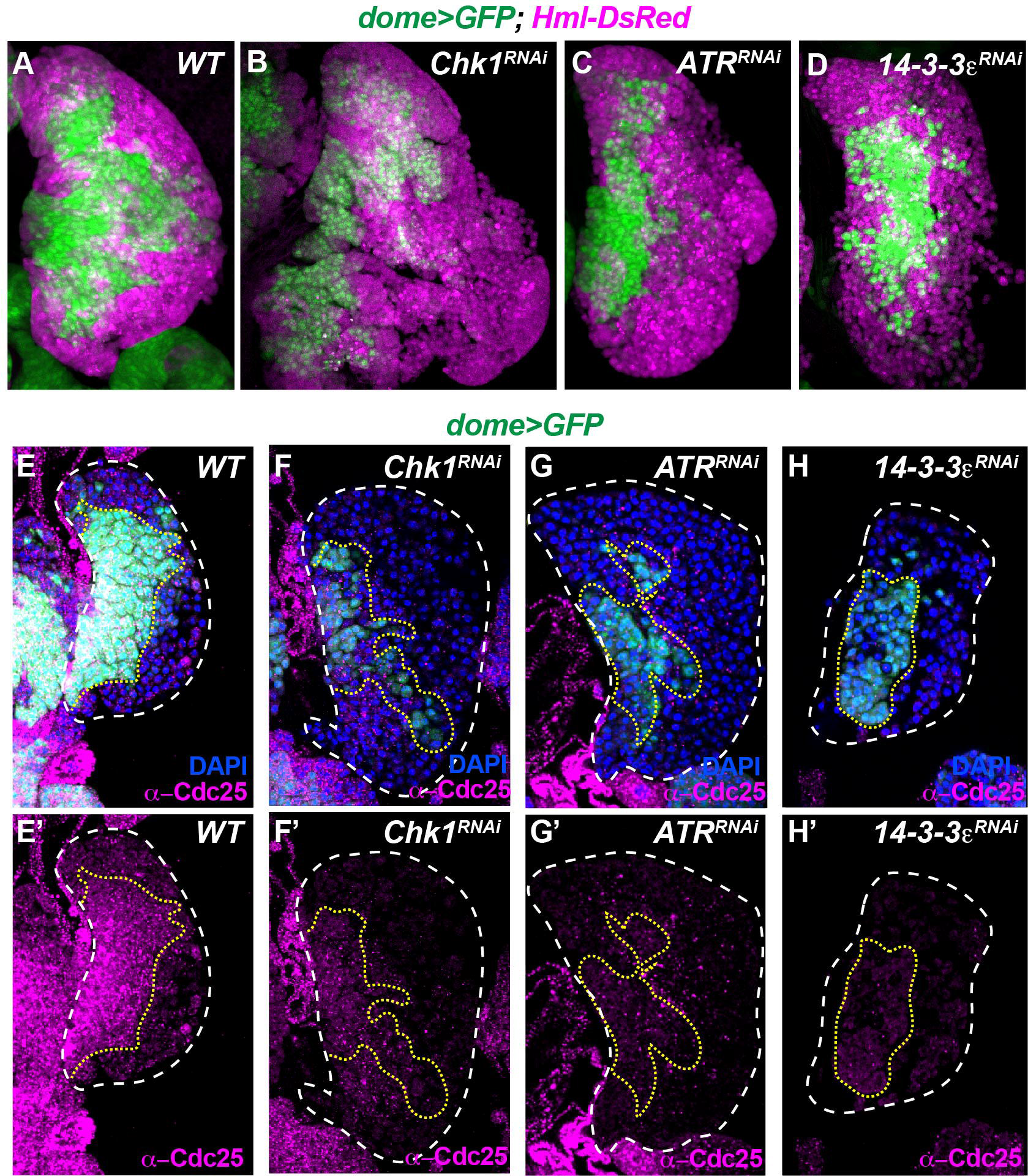
Mechanistic basis for Cdc25 sequestration that leads to a prolonged G2 phase. **(A-D)** Compared with wild type **(A)**, RNAi-mediated depletion of Chk1 **(B)**, ATR **(C)**, or 14-3-3ε **(D)** decreases the size of the progenitor population and increases the number of differentiated cells. Genotypes are as marked. GFP-positive progenitors (green), and *Hml*-positive cells (magenta) are shown for the middle third z-stack projections of representative lymph glands (LGs). (**E**-**H’**) Compared with wild type (**E**-**E’**), RNAi-mediated loss of Chk1 (**F**-**F’**), ATR (**G**-**G’**), or 14-3-3ε (**H**-**H’**), show a reduction in the size of the progenitor population (**F, G, H**) and reduced levels of Cdc25 protein staining **(F’, G’, H’**). The LGs are marked with *dome>GFP* (progenitors; green), DAPI (DNA; blue), and anti-Cdc25 staining (magenta). For clarity, the progenitor population and the outer boundaries of the lymph gland are demarcated by yellow-dotted and white-dashed lines, respectively. See also **Figure S2**.

### Wnt pathway in progenitor maintenance, Cdc25 sequestration, and cell cycle control

*Drosophila* 14-3-3ε (Par-5) was identified in oocyte polarization screens as binding to, and functioning together with, Par-1 (C-TAK1; MARK3) in several contexts ^65–67^. In unrelated studies, Par-1 was shown to bind Disheveled and to potentiate Wnt signaling^68, 69^. With these facts in mind, we investigated if the prolonged G2 phase in the hematopoietic progenitors is downstream of the Wnt pathway. Using appropriate, validated, RNAi constructs, we depleted the Wnt co-receptor LRP6/Arrow^70^ or the co-regulatory proteins, Ck1γ/Gish^71, 72^, Cyclin Y^73–75^, CDK14/Eip63E^19, 20^, and Par-1. Similar to that seen in Figure 2, we find that the loss of each of the above Wnt pathway-related proteins causes loss of progenitors with increased differentiation (Figure 3A-F; S3A-B), loss of Cdc25 (Figure 3G-L’), and a reduced percentage of G2 phase in the remaining progenitors (Figure S3C-D).

**Figure 3:**
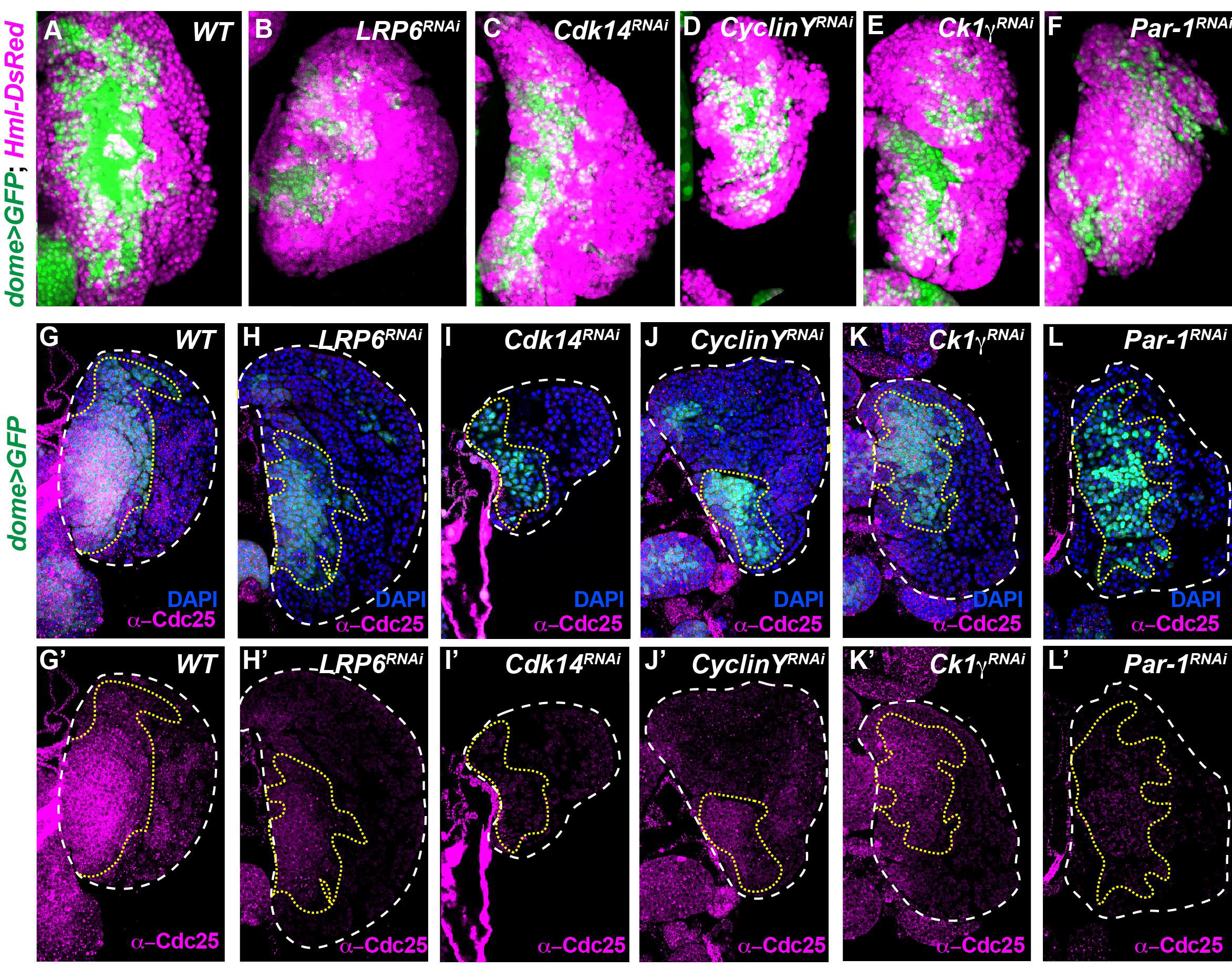
Wnt signaling pathway components control progenitor maintenance and Cdc25 levels. **(A-F)** Compared with wild type **(A)**, RNAi-mediated depletion of Wnt signaling components Arrow/LRP6 **(B)**, CDK14/Eip63E **(C)**, Cyclin Y **(D)**, CK1γ/Gish **(E)**, as well as the Wnt-associated kinase Par-1 **(F)** all cause a reduction in the number of progenitors and increased differentiation. Genotypes are as marked. Quantitation using nuclear markers is shown in **Figure S3**. **(G-L’)** Compared with WT lymph glands **(G,G’),** RNAi-mediated loss of Wnt signaling-related proteins Arrow/LRP6 **(H,H’)**, CDK14/Eip63E **(I,I’)**, Cyclin Y **(J,J’)**, CK1γ/Gish **(K,K’)**, and Par-1 **(L,L’)** all show a reduced progenitor population size (green; yellow-dotted line) and a major reduction in Cdc25 protein staining (magenta) in the remaining progenitors. The entire LG marked with DAPI/DNA is outlined with a white-dashed line. Markers: *dome>GFP* (green); DAPI/DNA (blue); anti-Cdc25 staining (magenta). See also **Figure S3**.

### Unique role of Wnt6 in cell cycle control and progenitor maintenance

The *Drosophila* genome contains seven Wnt proteins, each of which has corresponding mammalian orthologs^76^. We depleted each of the seven Wnts in the progenitor population and found that the loss of only a single ligand, Wnt6, gives rise to the entire set of the expected loss of function phenotypes including loss of progenitors, increased differentiation, loss of cytoplasmic Cdc25/String staining, and a decrease in the percentage of G2 in the remaining progenitors (Figure 4A-H). Knock-down of the remaining Wnt family ligands, Wnt1/Wg, Wnt2, Wnt3/5, Wnt4, Wnt8/D, or Wnt10 did not give rise to a similar loss of function phenotype (Figure S4A-H). A second independently generated *Wnt6-RNAi* line (Figure 4G, S4F) as well as a whole body Wnt6 loss of function mutant (*Wnt6^KO^*; Figure S4I-J), confirmed that Wnt6 controls progenitor maintenance. These results demonstrate that the pathway which sequesters Cdc25 in the cytoplasm, holding cells in G2 and thereby maintaining them as progenitors, is initiated by Wnt6. The similarities in the phenotypes of Wnt6 loss with that of the loss of stress and damage related proteins ATR/Chk1/14-3-3ε led us to believe that Wnt6 prolongs G2 during normal development by coopting several proteins associated with DNA damage-checkpoint control. In this model (Figure 4I), a Wnt6/LRP6 signal is transduced by Ck1γ, Cyclin Y, and CDK14.

**Figure 4:**
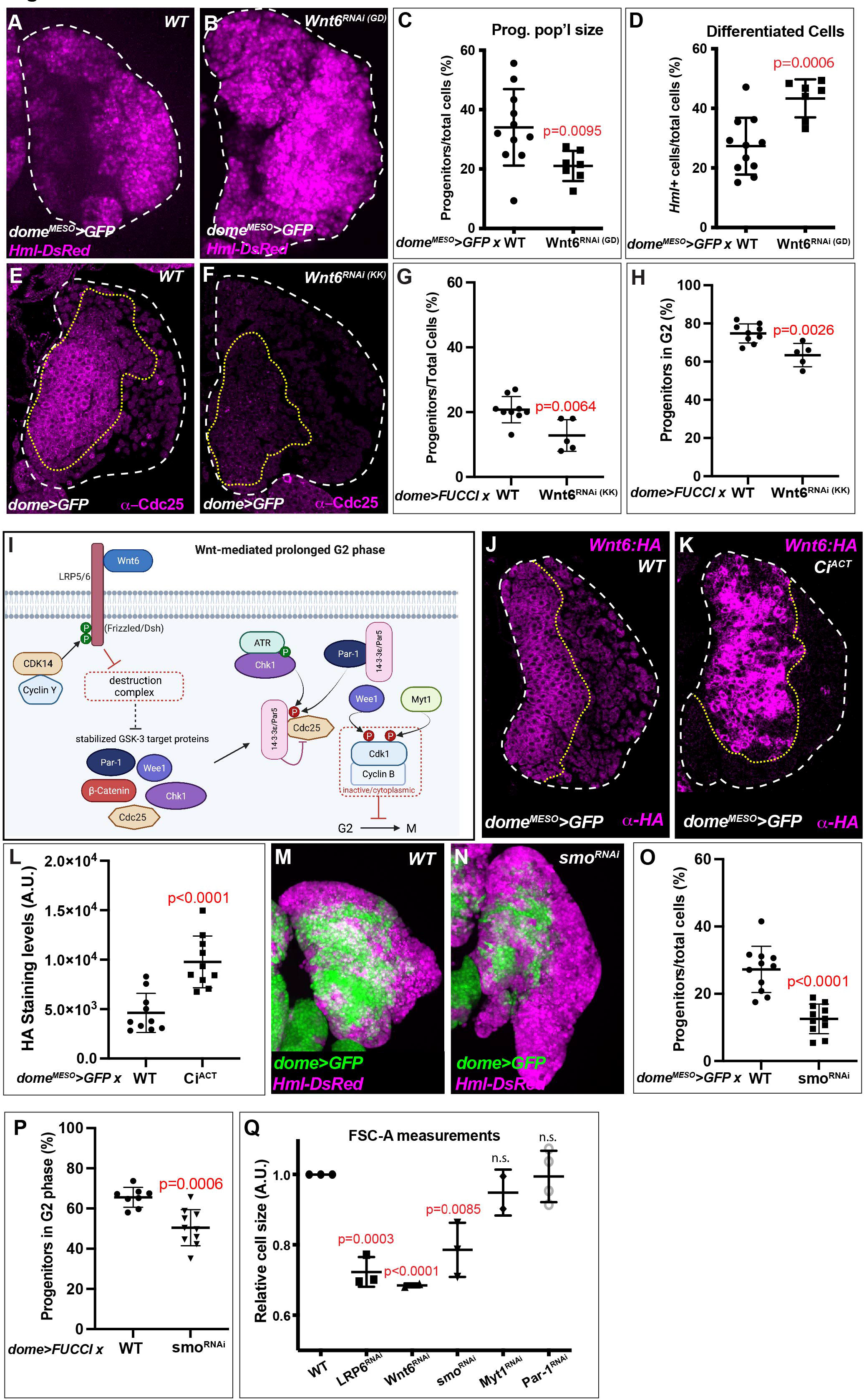
Wnt6 maintains progenitors in G2. **(A-C)** Compared to wild type (WT) **(A)**, *UAS-Wnt6-RNAi^GD^***(B)** in the progenitors (*dome^MESO^-GAL4*) increases the size of the differentiated cell population (magenta; *Hml-DsRed*). **(C-D)** Quantification of data in **(A-B)** shows a reduction in the size of the progenitor population **(C)** and an increase in the size of the differentiated cell population **(D)**. **(E-F)** Compared to wild-type lymph glands **(E)**, *UAS-Wnt6-RNAi^KK^* **(F)** in the progenitors (*dome>GFP*) results in a dramatic reduction in Cdc25 protein expression (magenta). **(G-H)** RNAi-mediated depletion of Wnt6 (*dome>FUCCI; UAS-Wnt6-RNAi^KK^)* results in a reduction in the size of the progenitor population **(G)** and in the percent of progenitors in G2 phase **(H)** compared with wild type at the same stage. **(I)** A model for a Wnt6 pathway-mediated signal that leads to cytoplasmic sequestration of Cdc25 and prolonged G2 phase. See text for details. **(J, K)** In wild-type **(J)**, progenitors show Wnt6:HA staining (magenta). Expression of a constitutively active Ci/GLI (*UAS-Ci^ACT^*; equivalent to Hh pathway activation) **(K)** in the progenitors (*dome^MESO^>GFP*) causes a dramatic increase in Wnt6:HA expression (magenta). **(L)** Quantification of Wnt6:HA staining levels from data in **(J)** and **(K)**. **(M-P)** Compared with wild type **(M)**, RNAi-mediated depletion of Smo **(N)** in the progenitors (green; *dome>GFP*) causes a reduction in their population **(O**; *dome^MESO^>GFP*), and a reduction in the fraction of progenitor cells in G2 (*dome>FUCCI*) **(P)**. **(Q)** Flow cytometric analysis of cell size (FSC-A) of the *dome^MESO^>GFP* population. A significant reduction in progenitor cell size is seen when Wnt6, its receptor LRP6, or Smo is depleted, but not upon the depletion of downstream proteins, Myt1 and Par-1. The GFP and DAPI channels in **(E-F)** and **(J-K)** are omitted for clarity but are used as the basis for demarcating the progenitor population (yellow dotted line) and the outer limits of the LG (white dashed line), respectively. See also **Figure S4**.

Further downstream they promote the cascading activity of Par-1^77, 78^, ATR, and Chk1 which are also associated with Wnt signaling^79^ and phenocopy loss of Wnt6. These downstream kinases are critical for Cdc25 phosphorylation (p-Cdc25) and its retention in the cytoplasm. While the upstream triggering mechanism is different between Wnt6 and radiation damage-induced block in G2, in either case, it is caused by the cytoplasmic retention of the p-Cdc25 /14-3-3ε complex.

### Wnt6, but not Wg, is highly expressed in the hematopoietic progenitors

The best studied Wnt ligand in *Drosophila* is Wingless (Wg; mammalian Wnt1). An antibody, Ab4D4, has been widely used to detect the Wg/Wnt1 protein (DSHB^80^). Based on staining with Ab4D4, we and others, in past studies, proposed that the lymph gland progenitors express Wg^81–83^. However, the current phenotypic analysis suggests that Wnt6, not Wg, functions in the lymph gland progenitors to maintain that population (Figure S4A-B, F). A well-controlled analysis of *Drosophila* maxillary palp development^84^ convincingly demonstrates that Ab4D4 cross-reacts with both Wg and Wnt6. The most parsimonious suggestion, based on our phenotypic analysis, would therefore be that Ab4D4 recognizes Wnt6 in the lymph gland. To substantiate this idea, we generated a Wnt6:HA direct protein reporter and compared its expression with that of the similarly produced Wg:GFP strain^85^. We find that Wg:GFP is not expressed in the lymph gland at this stage of development although its expression is strong in the disc epithelium (Figure S4K-M). In contrast, Wnt6:HA shows robust expression in the lymph gland progenitors (Figure 4J) as well as in the previously reported pattern for Wnt6 in the wing disc^84^ (Figure S4N,N’). As a control to validate the Wnt6:HA construct, we demonstrated that *Wnt6-RNAi*, but not *Wg-RNAi*, abrogates Wnt6:HA staining (Figure S4N-P’). The overall evidence conclusively demonstrates that at this stage of development Wnt6 functions in the hematopoietic progenitors.

### Hh signal from the PSC niche cells control Wnt6 transcription in the progenitors

Wnt6 protein is detected in the progenitor population, but how might its transcript be regulated in these cells? As Hedgehog (Hh) and Wg pathways show regulatory interactions in other tissues^76, 86^, we asked if Wnt6 might be related to Hh as well. It is known that the primary source of Hh in the lymph gland is the niche (PSC), and that the progenitors express Patched and Smoothened (Smo) as they are the recipients of the Hh signal^33, 38, 87, 88^.

We find that single-cell RNASeq analysis^41^ shows a strong positive correlation between *Wnt6* and *smo* transcripts in the MZ progenitors (Figure S4Q). Additionally, overactivation of the Hh pathway by progenitor-specific expression of Ci^ACT^ causes a substantial increase in Wnt6:HA but not in Wg:GFP levels (Figure 4J-L, S4M,R). Importantly, we find that RNAi-mediated depletion of Smo in the progenitors leads to increased differentiation, loss of progenitors, and a decreased percentage of progenitors in G2 phase (Figure 4M-P), similar to the phenotypes seen with loss of the Wnt6 pathway. We conclude that Wnt6 expression is specifically induced in the progenitors that receive a Hh signal from the niche. Since Wnt6 is a secreted molecule, presumably it can then be distributed to a broader population of MZ cells beyond the ones that directly receive Hh. The Wnt6 signal thus generated is responsible for progenitor maintenance and cell cycle control.

### Wnt6 and Hh signaling also control cell size of the progenitors

In the introduction section, we described the G2 phase growth-controlling pathway named Wnt-STOP and the concept that loss of this GSK3-dependent, β-catenin-independent signal gives rise to smaller cells^18^. We used flow cytometric analysis with FSC measurements to monitor cell size and found that RNAi-mediated loss of Wnt6 or LRP6 causes a significant decrease in cell size (Figure 4P), whereas loss of further downstream components such as Par-1 or Myt1 has no effect on cell size (Figure 4Q), even though all pathway members, including Wnt6, LRP6, Par-1, and Myt1 show strong evidence of cell-cycle control (Figures 4H, S3C,D, S2E, S4S). These results meet the expectations of Wnt-STOP signaling since only the components of the pathway upstream of the GSK3 kinase are expected to have G2-related function in cell growth^18^. Significantly, we find that in addition to its effect on the cell cycle, loss of Smo, belonging to the Hh pathway, also gives rise to a cell-size defect (Figure 4Q). This observation further bolsters the strong evidence that a Hh signal has an upstream role in the control of Wnt6 levels.

### Role of β-catenin in cell cycle regulation

Based on the Wnt6/LRP6 data on cell-cycle control, it is expected that β-catenin (*Drosophila* Arm), which is known to be stabilized upon the inhibition of the destruction complex, might also be required for progenitor maintenance. This, however, is not the case. Loss of β-catenin/Arm (*dome^MESO^-GAL4, UAS-β-catenin-RNAi*), in fact, shows quite the opposite phenotype from other Wnt6 pathway members. *β-catenin-RNAi* expressed in the progenitors causes an increase in the size of their population and a corresponding decrease in the number of differentiating cells (Figure 5A-D). Furthermore, unlike with loss of Wnt6 (Figure 4H) or LRP6 (Figure S3C), the loss of β-catenin does not decrease the fraction of progenitors that are maintained in G2 (Figure 5E). Since loss of β-catenin causes a mutant phenotype, this protein is functional in the tissue, but it is not responsible for holding progenitors in G2.

**Figure 5:**
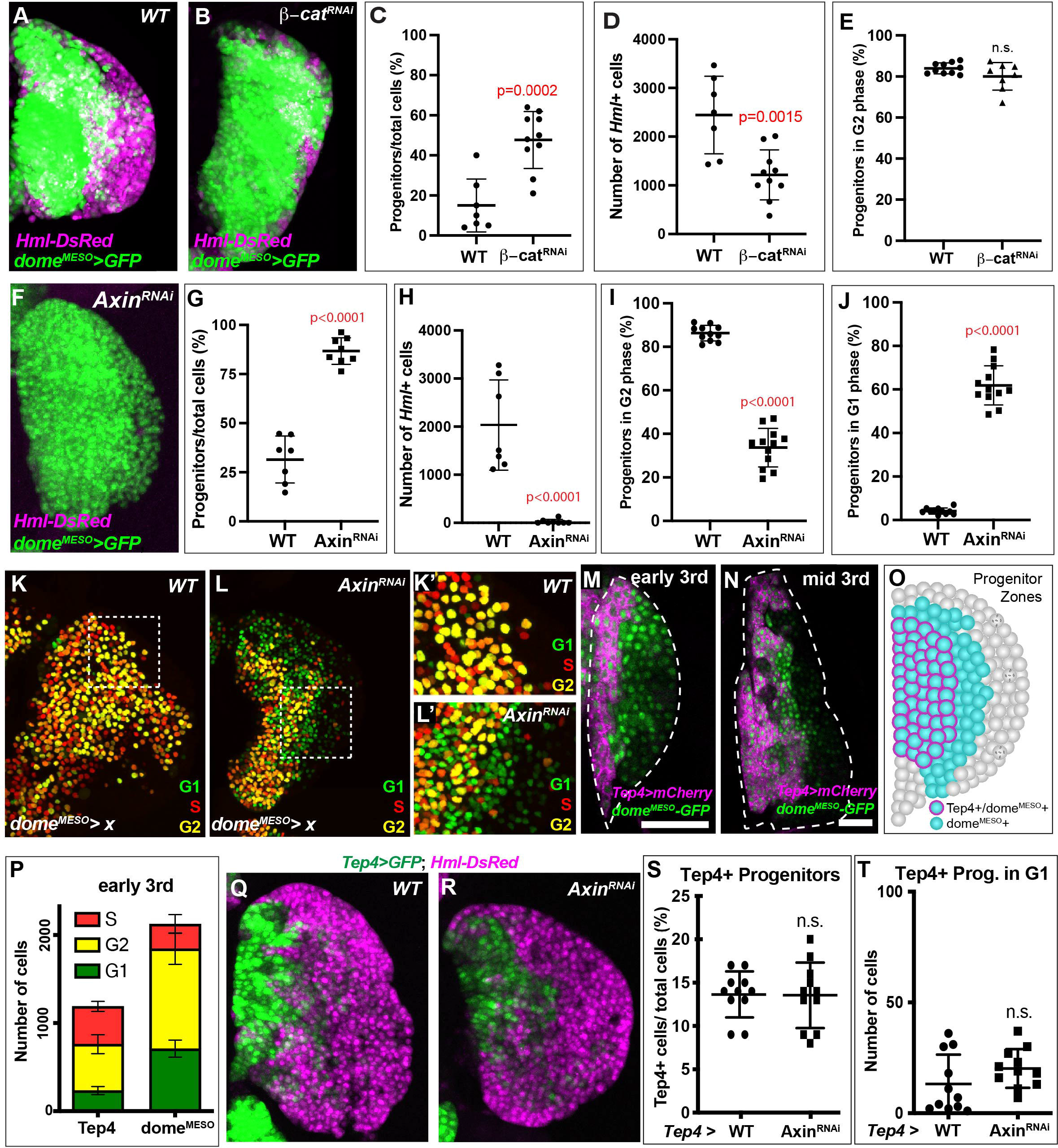
Role of β-catenin in cell cycle regulation. **(A-E)** Compared with wild type **(A)**, RNAi-mediated depletion of the β-catenin **(B)** results in an increased size of the progenitor population (green; *dome^MESO^>GFP*) and a significant reduction in differentiated cells (magenta; *Hml-DsRed*). **(C-D)** Quantification of data in **(A-B’)** shows that progenitor population size (expressed as the percentage of progenitors normalized to the total number of cells in each lymph gland) increases **(C)** while the total number of Hml-positive differentiating cells (IP or CZ) significantly decreases **(D)** upon RNAi-mediated depletion of β-catenin. In **(E)**, quantification of FUCCI data (*dome^MESO^>FUCCI*) shows that there is no change in the fraction of progenitors in G2 upon depletion of β-catenin. Note that all the phenotypes of β-catenin loss **(C-E)** are the opposite of that seen upon loss of the Wnt6 pathway components described in earlier Figures 3-4. **(F-J)** RNAi-mediated depletion of Axin, a component of the destruction complex, in the progenitors (*dome^MESO^>GFP*; *UAS-Axin^RNAi^*), equivalent to constitutive Wnt pathway activation, causes a complete loss of differentiated cells (magenta; *Hml-DsRed*) and increased numbers of progenitors (green; *dome^MESO^>GFP*) **(F)**. **(G-H)** The quantification of the results in **(F)** compared to WT. **(I-J)** Quantification of FUCCI data demonstrates a decrease in the fraction of the progenitor cells in G2 **(I)** with a huge increase in the fraction of progenitors in G1 **(J)** upon *Axin^RNAi^*. **(K-L’)** Location of progenitor cells in G1, S, and G2 (green, red, and yellow, respectively), as revealed by FUCCI analysis of wild type **(K,K’)** compared with Axin^RNAi^ **(L, L’)**, shows that the vast majority of additional G1 cells in Axin^RNAi^ **(L, L’)** are located at peripheral edge of the progenitor population (furthest from the dorsal vessel). **K’** and **L’** are higher magnification views of the region within the boxes in **K** and **L**, respectively. **(M-O)** Subpopulations of progenitors distinguished by live imaging of an early 3rd instar **(M)** or mid 3rd instar **(N)** lymph gland using the genetic combination: *Tep4-GAL4, UAS-mCherry; dome^MESO^-GFP* (mCherry shown in magenta and GFP in green). All progenitors express *dome*^MESO^ but only an inner subset of these cells, located closer to the heart than the rest, also express *Tep4*. This distinction is more apparent in earlier than in more mature larvae. These populations have been described by other investigators (citations in text), but their relative positions are best appreciated by live imaging. The *Tep4*-positive, *dome*-positive cells are referred to as “inner” or “core” progenitors. The *Tep4*-negative, *dome*-positive cells are the “outer” or “edge” progenitors. An idealized schematic is shown in **(O)**. **(P)** Cell cycle status of progenitor subpopulations with either *Tep4-GAL4* or *dome^MESO^-GAL4* driving *UAS-FUCCI* in early 3rd instar larvae. The total progenitor population (*dome^MESO^*-positive) is greater than the core progenitor population (*Tep4*-positive) and a majority of these additional cells are in G2 or G1 phases of the cell cycle. **(Q-T)** Compared with wild type **(Q)**, expression of Axin^RNAi^ **(R)** exclusively in the core progenitors (*Tep4>GFP*; green) has no effect on differentiation (*Hml-DsRed*; magenta). This is quantified in **(S)** using a nuclear marker (*Tep4>FUCCI*) with equivalent genotypes. Axin^RNAi^ also has no effect on the number of Tep4-positive progenitors in G1 phase **(T)**. This contrasts with **(F-H, J)**, where a similarly achieved activation of the Wnt pathway in all progenitors (*dome^MESO^-GAL4*) completely inhibits the formation of differentiated cells and greatly increases the number of *dome^MESO^-*positive progenitors in G1. See also **Figure S5**.

As a complement to the above loss-of-function approach, we enhanced β-catenin activity with a knock-down of Axin (*dome^MESO^-GAL4, UAS-Axin-RNAi*), which is a scaffolding protein and a key component of the destruction complex^89–91^. We find that the increased β-catenin activity in this genotype decreases differentiation and increases progenitor maintenance (Figure 5F-H). It also decreases the percentage of progenitors in G2 (Figure 5I). This is only in part due to the fact that the number of progenitors in G2 decreases, but also because the number in G1 increases by nearly 10-fold (Figure S5A-C). Ultimately this results in a rise in the percentage of cells in G1 from ∼5% in wild type to ∼50% when β-catenin levels are elevated (Figure 5J). Taken together, the loss and gain of function experiments suggest that β-catenin activity likely promotes exit from G2 and entry into G1, unlike the other Wnt6 components that are primarily involved in prolonging G2.

A closer look at the cell cycle status of *Axin-RNAi* progenitors within the lymph gland reveals that the additional G1 cells in this genotype are largely seen at the distal edge of the progenitor zone near the site of differentiation (Figure 5K-L’). We refer to these G1 cells, small in number in wild type (Figure S1G), and increased with excess β-catenin activity, as “edge progenitors”. The escape from G2 is localized to this distal edge and is not seen in the entire MZ, even though in these experiments the activity of β-catenin is altered throughout the MZ.

### Spatial regulation of the cell cycle

We took advantage of known differences in the expression patterns of two progenitor drivers (*Tep4-GAL4*^92, 93^ and *dome^MESO^-GAL4*^32, 92, 94, 95^) to better understand the spatial regulation of the cell cycle. While *dome^MESO^-GAL4* is ubiquitously active in all progenitors, *Tep4-GAL4* expression is confined to an inner subset of less mature *dome^MESO^*-expressing cells, which have been referred to as “core progenitors”, located closer to the dorsal vessel and farther away from the edge of the lymph gland^36–39^. The distinction in the expression of these two drivers is best appreciated in live imaging of the larva (Figure 5M-O). We find that the *dome^MESO^*-positive cells (“all progenitors’’) are separable into a *Tep4*-expressing subpopulation (“core progenitors”) and a *Tep4*-negative subset (“distal progenitors”). The extent of non-overlap between the two markers is greatest in the early 3rd instar, but by mid 3rd instar, this difference is over only a small number of cell diameters (Figure 5M-N). Combining these drivers with FUCCI allows us to also compare the cell cycle status of the progenitor subpopulations (Figure 5P; S5D,E). While both groups of progenitors (core and distal) are largely in G2, the core progenitors, particularly the cells close to the dorsal vessel, also show a significant number of cells in S phase (Figure S5F). On the other hand, the distal progenitors, while primarily in G2, also include cells in G1, particularly at their distal edge (Figure S1G). Thus, a progression of cell cycle is apparent, from an S/G2 state to a G2/G1 state in cells positioned from the inner to the outer reaches of the medullary zone.

Interestingly, the β-catenin-dependent canonical Wnt6 activity is distinct in the core vs distal progenitors. When β-catenin levels are raised (*Axin-RNAi*) in the core progenitors (with *Tep4-GAL4*), there is no significant effect on differentiation (Figure 5Q-S) or on the number of cells in G1 (Figure 5T). Whereas, as shown earlier (Figure 5F-H, S5A), an identical manipulation in all progenitors (using *dome^MESO^-GAL4*), causes a very efficient block in differentiation and increased numbers of progenitors in G1. We deduce that β-catenin-dependent Wnt6 pathway activity is spatially restricted to the edge of the medullary zone and only operates in distal progenitors where it helps mediate the release of progenitors from G2. In stark contrast, β-catenin-independent Wnt6 signaling is active in both core and distal progenitors since an increase in differentiation phenotype results when upstream members of this pathway (Wnt6, LRP6, or Par-1) or core cell cycle regulators (Wee1 or Myt1) are downregulated in either the core progenitors (*Tep4*-positive; Figure S5G-J) or in all progenitors (*dome^MESO^*-positive; Figure 4B; Figure 3B,F; Figure S2B,C). Together, these results imply that proper hematopoietic patterning requires the combined effect of a Wnt6 signal that is β-catenin-independent in all progenitors and a second, β-catenin-dependent signal that operates in a spatially restricted subset of distal progenitors.

### E-Cadherin and dual function of β-catenin in spatially restricted zones

E-Cadherin (E-cad) is an adhesion molecule known to be expressed in cells of the medullary zone^38, 82, 96, 97^. The core progenitors immuno-stain for both E-Cad (Figure S6A-B’) and β-catenin (Figure 6A-A’’’) at the cell surface, presumably representing the junctional complexes known to involve these proteins^98, 99^. The pan-β-catenin antibody used in this experiment (N2 7A1^100^) does not detect low levels of any endogenous nuclear β-catenin that might be present. The extensive network of E-Cad that holds the core progenitors together is sharply downregulated at the distal edge of the medullary zone (Figure S6A-A’’’). These E-Cad negative progenitors in the proximity of the distal edge are still *dome^MESO^*-positive but they are not part of the intermediate zone (Figure S6B,B’).

**Figure 6:**
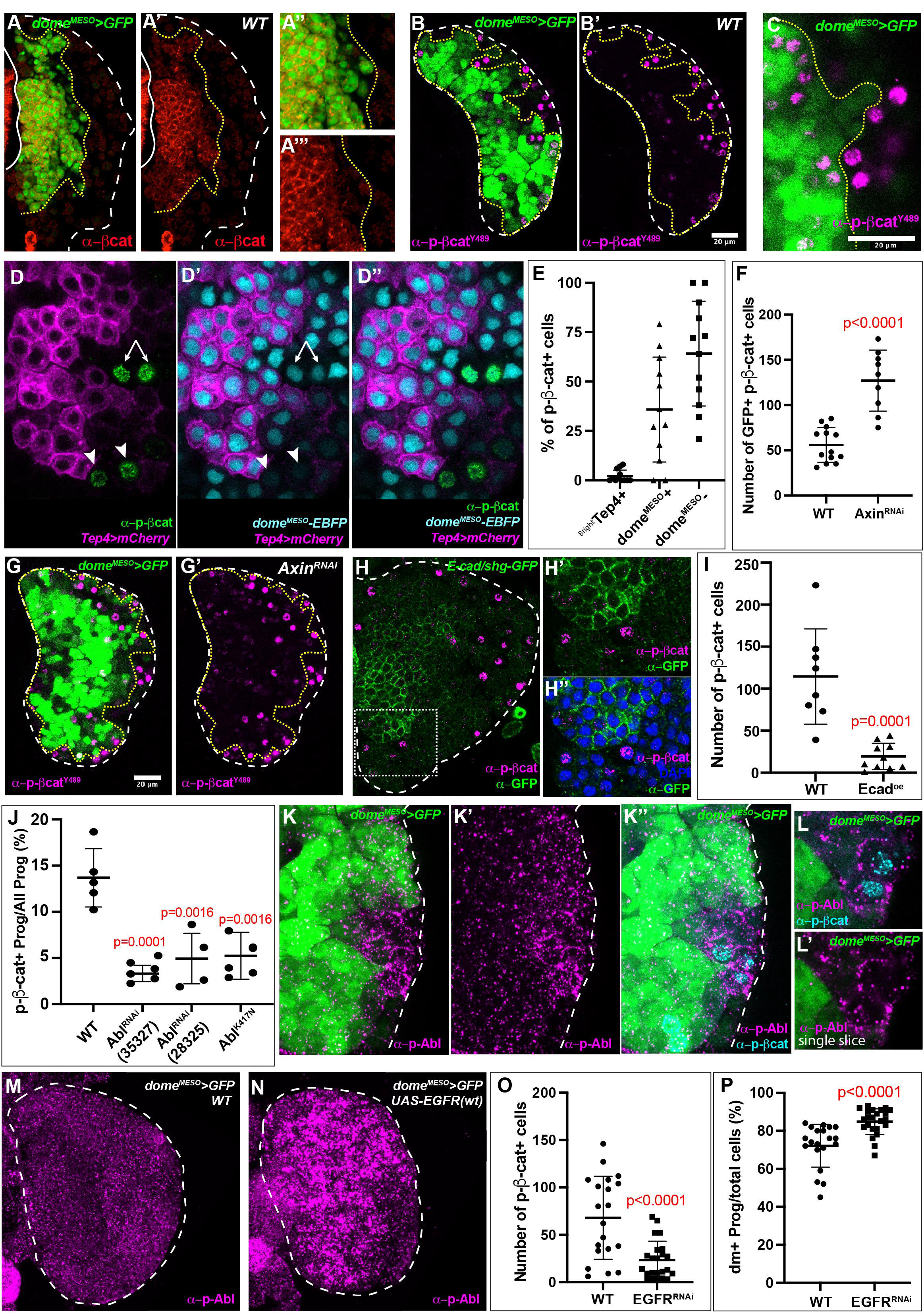
E-Cadherin and dual function of β-catenin in spatially restricted zones. **(A-A’’’)** Wild-type lymph glands with all progenitors marked (*dome^MESO^>GFP*; green) are stained with a pan-anti-β-catenin antibody (red) **(A)**. The antibody recognizes stabilized but non-nuclear protein that is seen in a honeycomb pattern at the cell surface of a subset of progenitors that are nearest to the dorsal vessel/heart **(A’)** but is absent from the cell surface of the outer progenitors closer to the distal edge. Higher magnification views **(A”, A’’’)** of the same lymph gland highlight the β-catenin expression (red) in only a subset of all progenitors (green). **(B-C)** phospho-specific anti-β-catenin antibody (anti-pY489-β-catenin) detects an active, nuclear form of β-catenin (magenta). **(B, B’)** When all progenitors are marked (*dome^MESO^>GFP*; green), nuclearly localized expression of activated β-catenin (magenta) is seen in distal (edge) progenitors with low expression of the progenitor marker (weak green), and also in their near neighbors, which are the most recent to have lost their progenitor markers (lack of green). No staining is detected in progenitors (green) nearest to the dorsal vessel/heart. **(C)** Higher magnification view of a similar lymph gland. **(D-D’’)** Wild-type lymph glands are doubly marked for all progenitors (*dome^MESO^-EBFP* direct fusion; cyan) and core (inner) progenitors (*Tep4-GAL4, UAS-mCherry*; magenta). Additionally, these glands are immunostained with anti-pY489-β-cat antibody (green). Nuclear pY489-β-catenin (green) is seen in a subset of progenitors that are *dome^MESO^*-positive (weakly cyan) but are *Tep4*-negative (arrows). Neighboring differentiating cells (arrowheads) that are negative for both *Tep4* (magenta) and *dome^MESO^* (cyan), also show evidence for active β-catenin expression. Core progenitors are *Tep4*-positive (magenta), and they do not express nuclear pY489-β-catenin. **(E)** Quantitative analysis of lymph glands in **(D-D’’)** shows that *Tep4*-positive core progenitors lack pY489-β-catenin. **(F)** Quantitative analysis demonstrates a highly significant increase in the number of pY489-β-catenin-positive progenitor cells upon RNAi-mediated loss of Axin. **(G-G’)** Representative examples of Axin-depleted lymph glands stained for active β-catenin, quantitated in **(F)**. Compare with WT **(B-B’**). **(H-H’’)** high Ecad-GFP expressing cells (green) do not express significant levels of nuclear pY489-β-cat (magenta). DAPI channel (blue) in **H”** is omitted in **(H,H’)** for clarity. **(I)** Over-expression of E-Cad in progenitors (Ecad^oe^; *UAS-DEFL*) causes a significant decrease in the number of pY489-β-cat-positive cells. **(J)** Loss of Abl in *dome^MESO^*-positive progenitors using independent loss of function constructs *UAS-Abl^RNAi^* (BDSC 35327), UAS-Abl^RNAi^ (BDSC 28325), or a dominant negative version of Abl, *UAS-Abl^K^*^417^*^N^*, each results in a significant decrease in the fraction of progenitors expressing nuclear pY489-β-catenin. **(K-L’)** pY412-Abl (magenta) and pY489-β-catenin (cyan) expression in wild-type lymph glands with progenitors marked (*dome^MESO^>GFP*; green). pY412-Abl levels increase in distal progenitors and differentiating cells. These cells also express pY489-β-catenin (cyan). Shown here in maximum intensity projections of several confocal slices **(K-K”)** or in magnified view of a single confocal slice **(L,L’)**. pY412-Abl staining (magenta) appears as puncta in the cytoplasm and near the surface of the cell while pY489-β-catenin staining (cyan) is in the nucleus. **(M,N)** Compared with wild type **(M)**, immunostaining intensity for pY412-Abl (magenta) greatly increases upon over-expression of wild-type EGFR **(N)**. **(O,P)** RNAi-mediated loss of EGFR in progenitors (*dome^MESO^>GFP*) results in a significant decrease in the number of pY489-β-cat-positive cells **(O)** and a significant increase in the normalized fraction of the progenitor population **(P)**. See also **Figure S6**.

While the inhibition of the destruction complex is the primary mechanism for stabilization of β-catenin^70, 101^, its subsequent phosphorylation at multiple sites dictates whether β-catenin localizes to the junctional complexes, stays in the cytoplasm, or is further modified to be stably transported to the nucleus where it can function as a transcription factor^22, 102^. In mammalian neuronal retina cells, one such terminal phosphorylation event generates phospho-Y489-β-catenin (pY489-β-cat for brevity), which is transported to, and is functional in, the nucleus^103^.

Furthermore, phosphorylation at this site decreases the affinity of β-catenin for cadherin complexes at the cell surface thus facilitating its release^103^. This phosphorylation site is conserved, although to our knowledge has not been studied in *Drosophila*. An antibody recognizing pY489-β-cat shows a spectacular staining pattern in the lymph gland, wherein pY489-β-cat is seen in the nuclei of a very restricted number of progenitors along the distal edge of the MZ and in their immediate differentiating neighbors of the transition zone (Figure 6B-C, S6C,C’). *Tep4*-positive core progenitors do not show any signs of nuclear pY489-β-cat expression (Figure 6D-E). Furthermore, forced overactivation of the Wnt signaling pathway downstream of the ligand (*dome^MESO^-GAL4, UAS-Axin-RNAi*) leads to an increase in the number of pY489-β-cat expressing cells (Figure 6F-G’, S6D), which are predominantly in G1 phase (Figure S6E).

When we directly compare E-cad localization and pY489-β-cat staining, we find that E-cad-positive progenitors do not colocalize with nuclear β-catenin (Figure 6H-H’’). Thus membrane-bound β-catenin, which is associated with E-cad and found in core progenitors, is non-overlapping with nuclear β-catenin, which is seen in the E-cad-negative distal progenitors. E-cad likely plays a crucial role in the regulation of β-catenin localization since over-expression of E-cad throughout the MZ (using *dome^MESO^-GAL4*) significantly reduces the number of pY489-β-cat-positive cells (Figure 6I). These results suggest a unique and novel relationship between Wnt6 signaling, E-cadherin function, pY489-β-cat nuclear localization, and the spatial regulation of the cell cycle.

### Abl controls nuclear β-catenin

In mammalian studies, the Y489 site has been shown to be recognized and phosphorylated by Abelson kinase (Abl)^103^. Such an Abl-related Y489 phosphorylation has not been investigated in *Drosophila*. We downregulated Abl activity in progenitors using two independent RNAi constructs and also using a dominant negative form of Abl^104^ and found that these genetic manipulations significantly decrease nuclear pY489-β-cat-positive progenitors (Figure 6J). This implies that Y489 phosphorylation and the consequent activation and nuclear translocation of β-catenin is indeed controlled by Abl. Activation of Abl, in turn, requires its own phosphorylation at the specific site, Y412^105–107^, and an available antibody against pY412-Abl identifies this active form in *Drosophila* ^108^. In a subset of progenitors largely at the edge of the MZ, pY412-Abl expression appears as punctate staining in the cytoplasm and at the cell surface (Figure 6K-L’, S6F-G’). Co-staining for pY412-Abl and pY489-β-catenin reveals that these two activated proteins localize to different compartments in the same cell (Figure 6K”,L). pY489-β-catenin is nuclear, whereas pY412-Abl staining appears as puncta in the cytoplasm and along the plasma membrane.

### Abl is activated by EGFR in a spatially restricted subset of progenitors

Abl activity can be controlled by a multitude of mechanisms^109, 110^. Prominent amongst these is its activation by the Receptor Tyrosine Kinase (RTK) EGFR^107, 111, 112^. We found that overexpression of wild-type EGFR notably increases the staining for active pY412-Abl (Figure 6M-N, S6H-I). This suggests that EGFR is involved in Abl phosphorylation and activation, possibly similar to what has been observed in mammalian tissue. Overexpression of wild-type EGFR normally does not have a phenotypic consequence in *Drosophila* as the limiting component of the pathway in *Drosophila* is the availability of the ligand^113, 114^. For the lymph gland, the data suggest that the amount of functional receptor is limited. This is an important concept and its implications for blood development are elaborated upon in the Discussion section. In order to bolster the results of the gain-of-function experiments, we knocked down EGFR levels with a pre-validated RNAi line which clearly decreases EGFR staining throughout the MZ (Figure S6J-K’). Consistent with our proposed phosphorylation cascade of EGFR/Abl/β-cat, we find that RNAi-mediated depletion of endogenous EGFR decreases overall numbers of pY489-β-cat-positive cells (Figure 6O). Together, these results support our model that EGFR activates Abl and Abl, in turn, phosphorylates β-catenin to promote nuclear translocation. Additionally, loss of EGFR also increases the size of the progenitor population (Figure 6P), suggesting that the EGFR signal plays a role in promoting differentiation.

### Linking cell cycle control with differentiation in distal progenitors

RTK signals have been proposed to have a regulatory role in differentiation in multiple tissues including the lymph gland^35, 115–118^. In such instances, they function by activating Pointed (Pnt/ETS1) as a downstream transcription factor^119^. When previously validated *pnt-RNAi* is expressed in all progenitors (*dome^MESO^-GAL4*) we observe a virtually complete block in differentiation^41^ (Figure 7A-C). However, when the expression of *pnt-RNAi* is limited to the core progenitors (*Tep4-GAL4*), there is no observed effect on differentiation (Figure 7D-F). This pattern of differentiation phenotypes is remarkably similar to that seen with enhanced β-catenin activity upon loss of Axin (Figures 5A,F,G, Q-S). In both instances, *pnt-RNAi* and *Axin-RNAi* function in the distal but not in the core progenitors. Also, in both cases, the total progenitor population is increased and that of differentiated cells is reduced. However, the composition in the numbers of cells in each phase of the cycle is different for these two genotypes in important ways that reveal differences in the underlying mechanisms. *Axin-RNAi* decreases the number of progenitors in S and G2 phases while increasing the number in G1 (Figure S5A-C; 7G). This suggests that β-catenin activity promotes exit from G2 and entry into G1, but limits progression through S phase and so does not promote proliferation in these cells. Instead, distal progenitors with unbridled β-catenin activity remain stuck in G1, unable to differentiate. In contrast, relative to wild type, *pnt-RNAi* causes an increase in the number of progenitors in all phases of the cell cycle with a nearly 3-fold increase in S phase, 2-fold increase in G2, and 8-fold increase in G1 (Figure 7H). This is an indication that differentiation of progenitors requires Pnt and normal Pnt activity limits the number of outer progenitors that proliferate. Without Pnt activity, the distal progenitors are unable to differentiate, and instead continue cycling, and are therefore populated by cells in all phases of the cell cycle.

**Figure 7:**
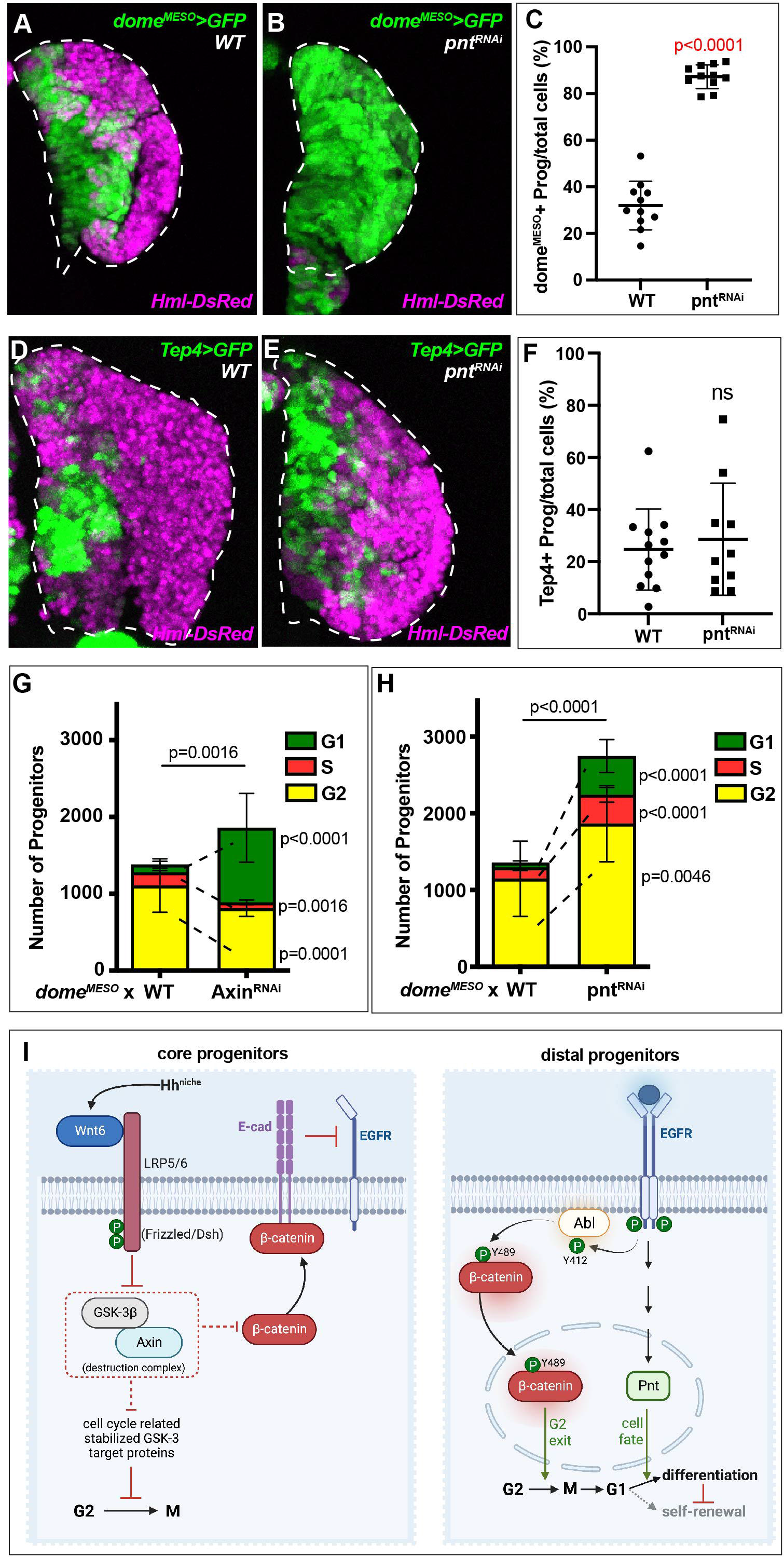
Linking cell cycle control with differentiation in distal progenitors. **(A-C)** Compared to WT **(A)**, essentially no differentiating cells (*Hml-DsRed*; magenta) are seen upon RNAi-mediated knockdown of Pnt **(B)** in all progenitors (*dome^MESO^>GFP*; green). **(C)** Quantification of similar results using *dome^MESO^>FUCCI* demonstrates a significantly increased fraction of progenitors upon Pnt downregulation. **(D-F)** Similar to the analysis in **(A-C)**, except with Pnt^RNAi^ driven in only the core progenitors (*Tep4>GFP*; green). In contrast with the dramatic loss of differentiated cells in **(B)**, loss of Pnt in the core progenitors has no effect on differentiation (*Hml-DsRed*; magenta) (magenta in **D,E** compared with **A,B**). **(F)** Quantitative analysis demonstrates that loss of Pnt in the core progenitors does not increase the size of the core progenitor population. **(G-H)** Quantification of the number of *dome^MESO^*-positive progenitors in each phase of the cell cycle (using *dome^MESO^>FUCCI*) seen upon gain of Wnt (Axin^RNAi^) **(G)** or loss of RTK pathway (Pnt^RNAi^) (**H)** compared to WT. **(G)** RNAi-mediated loss of Axin results in a significant increase in the number of the *dome^MESO^*-positive progenitors in G1 (green) and a significant decrease in G2 (yellow), and S (red). **(H)** RNAi-mediated loss of Pnt results in a significant increase in the number of the *dome^MESO^*-positive progenitors in all phases of the cell cycle including G2 (yellow), G1 (green), and S (red). **(I)** A mechanistic model for the interconnected regulation of progenitor maintenance and differentiation in core and distal progenitors. Core progenitors are held in G2 phase through Hh and Wnt6 signaling resulting in the stabilization of cell cycle-related proteins and β-catenin. The concerted activation of EGFR, Abl, and pY489-β-catenin in the distal progenitors induces G2 exit and differentiation. See Discussion for details. (Created with BioRender.com)

A synthesis of all the data presented allows us to propose a model for hematopoiesis (Figure 7I) in which dual functions of both Wnt and RTK pathways provide a link between cell cycle regulation and differentiation in the progenitors. Wnt6 is required for both blocking cells in G2 (β-catenin independent) and in releasing them from this block (β-catenin dependent) while EGFR is required for the activation of β-catenin (Pnt independent) and in promoting differentiation (Pnt dependent), with all EGFR activity localized to the distal progenitors. The key process that dictates such a direct coupling and thereby creates a homeostatic balance is the maintenance of the hematopoietic progenitors in an extended G2 phase (Figure 7I).

## Discussion

The multifunctional nature of Wnt6 signaling is at the core of our proposed model that explains the order of events that control the cell size, proliferation, maintenance, and differentiation of hematopoietic progenitors (Figure 7I). Combinatorial interactions between different forms of Wnt6 and tyrosine kinase signals ensure that only a subset of the progenitors proceed to differentiation while the rest are held in reserve. The rather elegant manner in which different modes of very few signals can generate multiple functional outputs ensures that maintenance, fate choice, proliferation, and cell size are intimately linked such that a change in one of these processes has a significant effect on the others. We strongly suspect that similar principles might underlie the regulation of mammalian multipotent progenitors, which are primary generators of cell-type diversity during native hematopoiesis^120, 121^. The components of the presented model are all conserved in evolution, but if this process is later found to be maintained in mammalian myeloid cells, based on the century of research in *Drosophila,* we should expect that while the overall logic of the model is likely to be conserved, the details of the exact pathways utilized and how they are combined, could vary somewhat. Also, the model (Figure 7I) explains all the data presented in this paper, but we fully expect that there are more components that belong to the scheme. For example, the role of free radicals such as ROS^122^ and NO (Cho *et al*., 2023) or any possible involvement of asymmetric cell division, will be critical to the model, and we hope to incorporate them in the near future.

As a brief synopsis, Hh expressed in the niche signals the nearest progenitors and causes *Wnt6* to be transcribed as its direct signaling target. Following its translation, modification, and secretion, Wnt6 initiates a signal that is transduced in all progenitors. This pathway inhibits the GSK3-dependent destruction complex, and many proteins that are GSK3 targets are thereby spared from degradation. The β-catenin-independent stabilization of a large number of such proteins constitutes the “Wnt-STOP” pathway^18^, which regulates the growth of the cell during the prolonged G2 phase, and a loss of pathway components upstream of GSK3, therefore, gives rise to reduced cell size. Additionally, this pathway also stabilizes key cell cycle regulators such as Cdc25, Par-1, Chk1, and Wee1^18, 21^. However, perhaps the most prominent β-catenin-independent function of the Wnt6 pathway, relevant to progenitor maintenance, is the cytoplasmic sequestration and inactivation of Cdc25 by 14-4-3ε. The phosphatase activity of nuclear Cdc25, critical for the activation of the nuclear Cyclin B/CDK1 complex, is the primary facilitating event in the G2-M transition of a mitotic cell^52, 53^. Cytoplasmic sequestration of Cdc25 maintains the progenitors in a prolonged G2 phase until this block is lifted only in the most mature subset of progenitors.

Stabilized nuclear β-catenin, a hallmark of “canonical Wnt signaling”, also functions in this system but rather than lengthening G2, it is essential to release only the most mature progenitors from G2, allowing a round of mitosis to generate daughters in G1 that are then competent to choose whether to self-renew or differentiate. Although β-catenin degradation is inhibited in all progenitors, its nuclear transcriptional activity is spatially restricted to only a small number of cells in which it is phosphorylated by Abl, which is activated by EGFR. Interestingly, the expression of EGFR is not restricted to this small subset of the most mature progenitors. Rather, it is the activity of EGFR that is restricted by a combination of mechanisms. First, in their elegant analysis of the system, Cho *et al*. (2023) have established how the EGFR protein is inactivated by nitrosylation and is unable to exit the ER in all but the most mature progenitors.

And second, any EGFR escaping this process will also be inactive in the high E-Cad environment of the core progenitors^123–126^. In *Drosophila*, spatially restricted EGFR activity is usually a result of limited expression of the ligand^113, 114^. This is not the case for the lymph gland as both this study as well as Cho *et al*., show that it is the availability of functional receptor that is limiting. This mechanism is the norm in mammalian systems^127, 128^ and perhaps reflects the uniquely conserved strategies adopted for hematopoietic progenitor maintenance. In summary, active EGFR activates Abl, which in turn facilitates a specific phosphorylation event that reduces β-catenin’s affinity for the junctional complex and promotes its transfer to the nucleus^103^, where its transcriptional activity enables only the distal progenitors to escape their G2 block and proceed through mitosis with the daughter cells in G1 awaiting differentiation signals.

The direct coupling between cell cycle and differentiation is made closer still because the EGFR pathway, like Wnt6, also plays multiple roles. Its transcription-independent arm phosphorylates Abl and is ultimately responsible, in the context of Wnt6 signaling, for release of the progenitor from the G2 phase. But its canonical Ras/Raf/Pnt arm produces a signal that the emerging cell in G1 perceives as a differentiation cue. A majority of these cells follow a differentiation path and very few self-renew. The latter process becomes prominent only when differentiation is blocked, such as when Pnt is depleted in the distal progenitors. That EGFR functions in both G2 exit and in differentiation is well supported by our data, but this does not rule out that another RTK, such as the two FGFRs (Htl^117^ and Btl^35^), might also be involved in these processes in a partially redundant manner. Additional such Pnt-dependent activities, if present, will not fundamentally alter the model.

The ability to maintain a reserve number of progenitors while also allowing a small number of them to escape their *status quo* and differentiate along a spatial boundary within the confines of a hematopoietic organ requires an intimate spatio-temporal link between the mechanisms that control cell cycle and those that favor differentiation. What is so elegant about how hematopoiesis achieves this is the combinatorial use of various forms of a very small number of signaling systems. This is particularly valuable since a large majority of the progenitors that are maintained are not in direct contact with their niche, and this contrasts with the very small populations of traditionally defined stem cells that are kept quiescent by their proximity to neighboring niche cells^129^. Unlike transient amplifying cells, the hematopoietic progenitors become “quiescent” for a significant proportion of the time that they are maintained, but they utilize a very different strategy than what stem cells employ. They achieve their period of quiescence by lengthening G2, rather than G1/G0.

This strategy might be considered wasteful as G2 arrest implies that the entire population is made up of cells with replicated DNA (4N) over a long period of time risking damage and yet unable to perceive signals that would directly result in differentiation. We speculate that an undeniable advantage that this affords to a progenitor is that it can very rapidly respond to environmental stress and infection and produce the repertoire of immune cells akin to those of the myeloid lineage without first having to replicate its DNA in an S-phase. In fact, past work has established that induction of such environmental stress causes a rapid deployment of immune cells rather than significant increases in the progenitor population. Furthermore, work from our lab and others have found that the same signaling pathways employed here to maintain homeostasis are also co-opted and function in response to nutritional stress^81^ (Wnt) and immune challenge^130^ (EGFR). Such dual signaling mechanisms allow for maintaining a homeostatic balance between hematopoietic progenitors and their differentiated progeny and also provide a means to tune that balance in response to stress, a key feature of the hematopoietic system across species.

Several aspects of *Drosophila* hematopoietic development are conserved in humans. Of the two major branches of mammalian hematopoiesis, it is the one leading to the myeloid lineage that is ancient in evolution, with cell types similar to macrophages conserved in very primitive invertebrates. The notion that generation of myeloid cell type diversity, important for sensing stress, inflammation, and innate immune challenge, is more a function of the multipotent progenitors than of the rarely dividing HSCs is gathering strength in recent mammalian studies^131^. Whether this means that, in humans, a large progenitor pool is maintained *in vivo* and utilizes a controlled mechanism similar to that we see in this study is a matter of speculation. But given the past history of conserved mechanisms between *Drosophila* and mammals, it seems a likely possibility worthwhile to investigate in future studies.

## Acknowledgments

We gratefully acknowledge FlyBase, the Developmental Studies Hybridoma Bank (DSHB), the Bloomington *Drosophila* Stock Center, the Vienna *Drosophila* Resource Center, the Kyoto Stock Center, and the fly community for reagents. We acknowledge the help of the Broad Stem Cell Research Center (BSCRC), the MCDB/BSCRC Core Facility in Microscopy, and the BSCRC core in Flow Cytometry. We thank undergraduate research scholars Khoi Luc and Stephanie Bottomley for their contributions. U.B. is supported by National Institutes of Health grants R01HL067395 and R01CA217608; L.M.G. by Ruth L. Kirschstein Institutional National Research Service Award number T32CA009056 and the National Institute of Diabetes and Digestive and Kidney Diseases of the National Institutes of Health under Award number K01DK132488; J.R.G. by Ruth L. Kirschstein National Research Service Award number T32HL69766 and UPLIFT (UCLA Postdocs’ Longitudinal Investment in Faculty) Award number K12GM106996.

## Author Contributions

Conception and writing of the manuscript: L.M.G. and U.B. Experimental design: L.M.G., J.R.G., and B.C.M. Data generation, analysis, and imaging: L.M.G., J.R.G., B.C.M., S.B., C.C.S., R.T., and T.B

## Methods

### Drosophila strains

The *Drosophila* lines used in this study were from our lab stocks, the Bloomington *Drosophila* Stock Center (BDSC), the Vienna Drosophila Resource Center (VDRC), the National Institute of Genetics (NIG-Fly), or kind gifts from other labs as indicated in the associated chart. Briefly, lines from our lab stocks include: *dome^MESO^>GFP Hml*^Δ^*-DsRed*, *dome^MESO^-EBFP2, CHIZ-GAL4 UAS-mGFP* (IZ-specific *GAL4*^42^), *Tep4>GFP Hml*^Δ^*-DsRed*, *domeless>GFP*; *Hml-DsRed*, and *dome^MESO^*-GAL4 UAS-FlyFUCCI. Wnt6:HA was generated in this study through Recombinase-Mediated Cassette Exchange (RMCE) of the GAL4 sequence with an HA tag in the Wnt6-CRIMIC-TG4 line^132^ (RMCE performed by WellGenetics, Inc). The potency of the RNAi lines used have been confirmed and multiple RNAi lines were used to validate findings.

### Lymph gland dissection, immunostaining, and imaging

Crosses were performed at 25°C and the resulting eggs were shifted to 29°C to maximize GAL4/UAS expression. Due to differences in the growth rates of individual stocks and crosses, larvae were developmentally staged according to age and size. Lymph gland size has been previously shown to be highly reproducible and stage-specific^42^. 1^st^ instar larvae were chosen from vials ∼24-48 hours after egg lay (hAEL), early 2^nd^ instar larvae and late 2^nd^ instar larvae were both chosen from vials ∼48-72 hAEL and divided by the number of cells in the lymph gland into early 2^nd^ (below 750) or late 2^nd^ (above 750). Early 3^rd^ and mid 3^rd^ instar larvae were both chosen from vials ∼72-96 hAEL and divided based on the size of the larva. Larvae that were found wandering up the size of the vials (∼96-120 hAEL) and out of the food were designated wandering 3^rd^ instar (w3rd) larvae.

Lymph glands were dissected and processed as previously described^32^. Unless indicated otherwise in the figure legend, all stainings were performed on lymph glands from wandering 3rd instar larvae. For Wnt stainings (Wg:GFP with anti-GFP, Wnt6:HA with anti-HA, and Ab4D4), 10% NGS in PBS was used for blocking and antibody incubations. E-cadherin staining was done as previously described^97^. For all other stainings, lymph glands were dissected into cold PBS and then fixed in 4% formaldehyde in PBS at room temp for 25 minutes. After fixation, tissues were washed four times in PBS with 0.3% Triton X-100 (PBST) for 10 minutes each, blocked in 10% normal goat serum in PBST (blocking solution) for 30 minutes, followed by incubation with primary antibodies in blocking solution overnight at 4°C. The following day tissues were washed four times in PBST for 10 minutes each, followed by incubation with secondary antibodies for 2-4 hours at room temperature. Samples were washed four times in PBST, with DAPI (1:500, Invitrogen) or ToPro-3 (1:1000, Invitrogen) added to the third wash to stain nuclei, and then placed into VectaShield mounting medium (Vector Laboratories) and mounted on glass slides. Lymph glands were imaged using a Zeiss LSM 880 confocal microscope. All microscopy data are representative images from a total of approximately 10 biological replicates (n) in most cases. For each n, z-stacks were imaged, processed, and analyzed using ImageJ or Imaris software. For cell type and cell cycle quantifications, Imaris software was used to reconstruct a 3D volume from a z-stack and the spots feature was used to quantify individual cell types based on their fluorescence intensity values. ImageJ or Imaris was also used to assess protein staining levels. For all quantification graphs, the mean and standard deviation are shown and significance was calculated by unpaired t-test. All statistics were performed using Prism software (GraphPad) and p-values are shown in charts or figure legends as indicated. For space considerations, in some graphs p-values are represented using asterisks as follows: n.s. if p>0.05; * if p≤0.05; ** if p≤0.01; *** if p≤0.001; **** if p≤0.0001.

The primary antibodies used are as follows: mouse anti-Wg (1:100; DSHB Ab4D4-c), rat anti-HA (1:100; Sigma clone 3F10), mouse anti-β-catenin (1:10; DSHB N2 7A1 Armadillo-s), rabbit anti-Cdc25/String (1:500; kind gift of Dr. Eric Weischaus), guinea pig anti-Cdc25/String (1:500; kind gift of Dr. Yukiko Yamashita), mouse anti-Cyclin B (1:10; DSHB F2F4-s), mouse anti-Cyclin A (1:5; DSHB A12-s), mouse anti-phospho-Y489-β-catenin (1:200; DSHB PY489-B-catenin-c), rat anti-Ecad (1:50; DSHB DCAD2), rabbit anti-GFP (1:100; Invitrogen A-11122), rabbit anti-Pxn (1:500; kind gift of Dr. Jiwon Shim), mouse anti-dEGFR (1:100; Sigma C-274). Primary antibodies were detected with secondary antibodies conjugated to Cy3 (1:100; Jackson ImmunoResearch Laboratories), Alexa Fluor 633 (1:100), Alexa Fluor 555 (1:200), or Alexa Fluor 488 (1:200) (Invitrogen).

### Live Imaging

Lymph glands were observed in their most native state by visualizing through the live larvae. Larvae were first transferred from vials into water-filled glass dishes on ice. The cold-shocked larvae were then placed on glass slides with 5 drops of water and oriented perpendicular to the length of the glass. A glass coverslip was then carefully placed over the larvae and excess water was removed through capillary action to increase the pressure on the larva, which helped immobilize it. The size of the coverslip varied with the size of the larvae (1^st^ and 2^nd^ instar 22 mm x 22 mm, early 3^rd^ instar 22 mm x 40 mm, and wandering 3^rd^ instar 22 mm x 50 mm) in order to exert the correct amount of pressure without killing the larva. The prepared slide was viewed using a laser scanning confocal microscope (Zeiss LSM700). The coverslip was then shifted using a small wooden baton to rotate the larvae until the lymph gland was in the best position for imaging. The entire process (including imaging) was performed in less than 15 minutes to minimize stress on the larva.

### Flow cytometry

For DNA content analysis, dissociated lymph gland cells were fixed in 1 mL of 1% formaldehyde solution in PBS after being dissociated using the dissociation protocol described in Girard *et al*.^41^. Cells were incubated in fixative in low binding tubes for 30 minutes at 4°C on a shaker, then were spun down and washed with PBS. Fixed cells were resuspended in a solution of PBS containing NucBlue™ Live ReadyProbes™ Reagent (Hoechst 33342) and incubated at room temperature on a shaker for 30 minutes. Cells were transferred to 5 mL polystyrene tubes for flow cytometry analysis on the BD LSRII. Cells were gated to exclude doublets using FSC-H vs FSC-W and SSC-H vs SSC-W comparisons. Relative cell size was determined by comparing FSC-A measurements. The progenitor (GFP-positive) population was gated and average FSC-A in each mutant genotype was calculated relative to FSC-A measurements in the corresponding WT genotype performed on the same day with the same parameters.

### Single Cell RNA Sequencing Analysis

See Girard *et al*.^41^

---

*Drosophila* Stocks:

| Genotype | Source |
| --- | --- |
| w<sup>1118</sup> | Banerjee Lab |
| domeMESO>FUCCI | Banerjee Lab |
| domeMESO>GFP, Hml-DsRed | Banerjee Lab |
| UAS-Dcr2; domeMESO>GFP, Hml-DsRed | Banerjee Lab |
| domeMESO-GFP/CyO | Banerjee Lab |
| UAS-Dcr2; domeMESO>GFP | Banerjee Lab |
| domeMESO>GFP | Banerjee Lab |
| domeMESO>GFP-Tubulin | Banerjee Lab |
| domeMESO-EBFP | Banerjee Lab |
| Wnt6:HA; domeMESO>GFP | Banerjee Lab |
| Wg:GFP; domeMESO-GAL4 | Banerjee Lab |
| domeless>GFP | Banerjee Lab |
| domeless-GAL4; ; UAS-FUCCI | Banerjee Lab |
| domeless>FUCCI | Banerjee Lab |
| domeless>GFP, Hml-DsRed | Banerjee Lab |
| Tep4>mCherry/CyO | Banerjee Lab |
| Tep4-GAL4; UAS-FUCCI | Banerjee Lab |
| Tep4>GFP, Hml-DsRed | Banerjee Lab |
| Tep4>mCherry | Banerjee Lab |
| CHIZ>mGFP | Banerjee Lab |
| Wg:GFP | Fillip Port |
| Wnt6:HA; dpp>GFP | Banerjee Lab |
| Nup98-GFP | Banerjee Lab |
| Shg-GFP (trap) | BDSC: 60584 |
| UAS-Cdc25 or UAS-Stg | BDSC: 4777 |
| UAS-Lgl-mCherry | Knoblich Lab |
| UAS-Pon-GFP | Jan Lab |
| UAS-DEFL (Ecad(wt)-GFP) | Yamashita Lab |
| Chk1-RNAi | VDRC: v110076-KK |
| Chk1-RNAi | VDRC: v12680-GD |
| ATR-RNAi | VDRC: v103624-KK |
| 14-3-3epsilon-RNAi | BDSC: 31497 |
| 14-3-3epsilon-RNAi | BDSC: 35441 |
| 14-3-3epsilon-RNAi | VDRC: 108129-KK |
| ATR-RNAi | VDRC: 103624-KK |
| Myt1-RNAi | VDRC: 105157-KK |
| Wee1-RNAi | VDRC: 106329-KK |
| LRP6-RNAi or arrow-RNAi | BDSC: 53342 |
| LRP6-RNAi or arrow-RNAi | VDRC: 6707-GD |
| Cdk14-RNAi or L63-RNAi | VDRC: 47860-GD |
| CycY-RNAi or CyclinY-RNAi | BDSC: 34009 |
| CycY-RNAi or CyclinY-RNAi | VDRC: 107010-KK |
| CK1gamma-RNAi or gish-RNAi | BDSC: 35138 |
| Par-1-RNAi or Par1-RNAi | BDSC: 35342 |
| Wnt6-RNAi | VDRC: 27610-GD |
| Wnt6-RNAi | VDRC: 104020-KK |
| Wnt6<sup>KO</sup> | Teleman Lab |
| UAS-Ci<sup>ACT</sup> or UAS-HA.ci.m1-3*103 | BDSC: 32571 |
| Smo-RNAi | BDSC: 24472 |
| Smo-RNAi | Banerjee Lab: 774 |
| Wg-RNAi or Wnt1-RNAi | VDRC: 13351-GD |
| Wg-RNAi or Wnt1-RNAi | VDRC: 13352-GD |
| Wg-RNAi | VDRC: 39676-GD |
| Wnt2-RNAi | VDRC: 38079-GD |
| Wnt3-RNAi or Wnt5-RNAi (Wnt3/5-RNAi) | VDRC: 32257-GD |
| Wnt4-RNAi | VDRC: 38010-GD |
| Wnt8-RNAi or WntD-RNAi (Wnt8/D-RNAi) | VDRC: 107727-KK |
| Wnt10-RNAi | VDRC: 100867-KK |
| Arm-RNAi or Beta-catenin-RNAi | BDSC: 31305 |
| Axin-RNAi | BDSC: 31705 |
| Abl-RNAi | BDSC: 35327 |
| Abl-RNAi | BDSC: 28325 |
| UAS-Abl(K417N) or UAS-Abl(DN) | BDSC: 8566 |
| UAS-EGFR.2(wt) | BDSC: 9535 |
| EGFR-RNAi | VDRC: 43267-GD |

## Supplemental Figures

**Movie 1:** Z-stack time-lapse live imaging of a w3rd instar lymph gland with progenitors expressing *dome^MESO^>FUCCI*. The transmitted light channel (gray) is omitted from the images on the right for clarity and a white line outlining the periphery of the lymph gland is added. This movie illustrates the preponderance of progenitors in G2 phase (yellow) at this stage of development.

**Figure S1:**
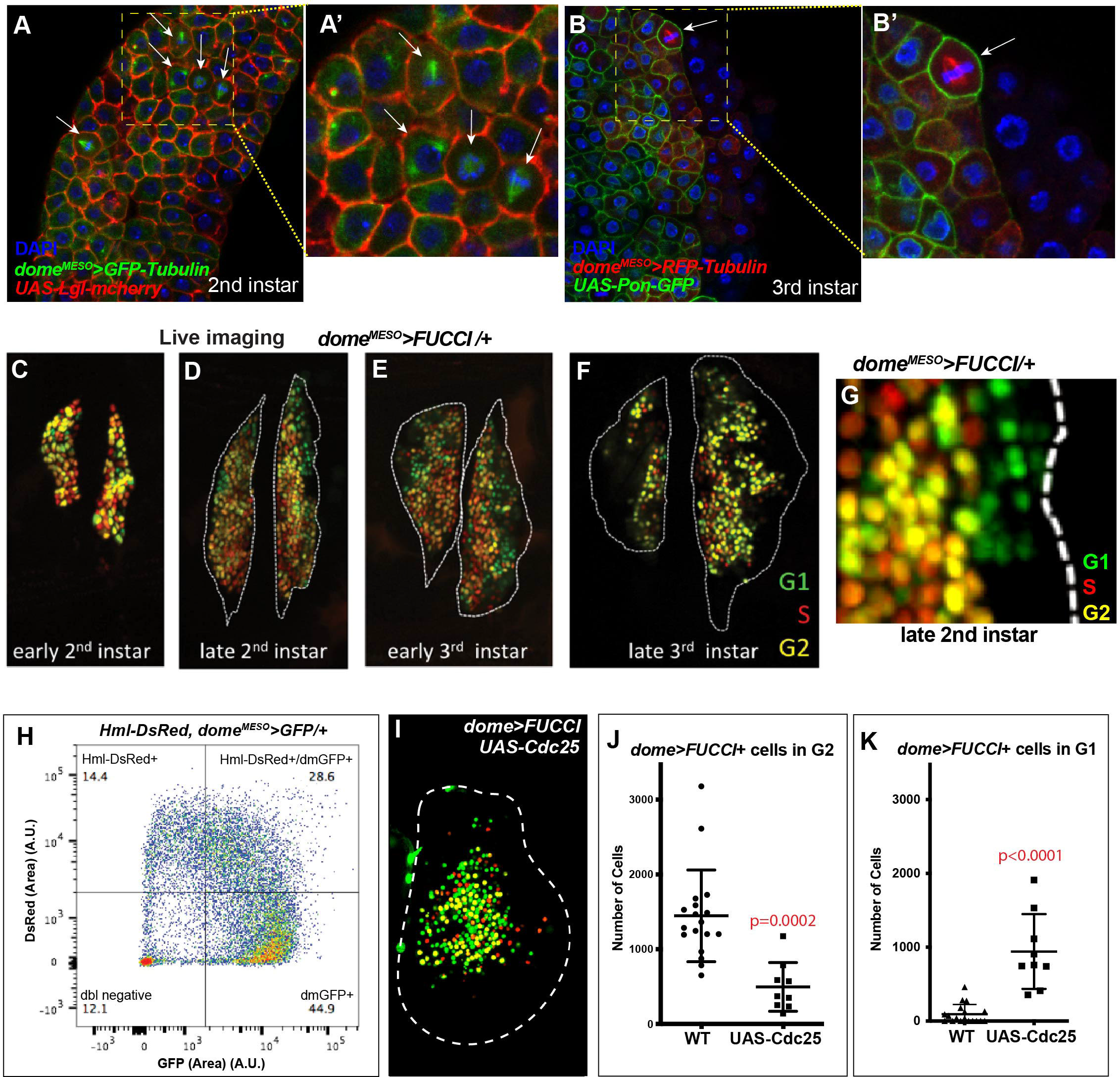
Hematopoietic progenitors are held in G2 phase of the cell cycle. **(A-A’)** Representative image of a lymph gland (LG) from a 2nd instar larva. Mitotic cells (arrows; identified by the morphology of the tubulin network) can be found throughout the progenitor population. **(A’)** higher magnification view of the region outlined with a dashed line in **(A).** Genotype: *dome^MESO^-GAL4; UAS-GFP-Tubulin; UAS-Lgl-mCherry*. **(B-B’)** Representative image of a LG from a 3rd instar larva. Mitotic cells (arrows; identified by the morphology of the tubulin network) are largely found at the distal edge of the progenitor population near the site of differentiation (identified by RFP/GFP-negative cells in which *dome^MESO^-GAL4* has turned off and therefore these cells only show DAPI staining). **(B’)** higher magnification view of the region outlined with a dashed line in **(B).** Genotype: *dome^MESO^-GAL4; UAS-RFP-Tubulin; UAS-Pon-GFP*. **(C-F)** Live imaging of LGs expressing *dome^MESO^>FUCCI* in intact larvae at different stages of development. A similar cell cycle profile is seen in live imaging as that seen in fixed and stained tissues (Figure 1B-F). Live imaging better maintains the 3D structure of the LG. The periphery of the LG is indicated with a white dashed line as determined by transmitted light imaging. **(G)** Live imaging of a LG in late 2nd instar expressing *dome^MESO^>FUCCI*. G1 cells (green) are largely restricted to the edge of the progenitors near differentiating cells that have turned off *dome^MESO^-GAL4* and therefore appear dark. **(H)** Scatter plot of the GFP and DsRed fluorescence levels observed in dissociated cells from *Hml-DsRed, dome^MESO^>GFP* LGs. GFP-positive progenitors (lower right), DsRed-positive/GFP-positive intermediate progenitors (upper right), and *Hml-DsRed*-positive differentiating cells (upper left) can be separated into quadrants and gated. The DNA content of each of these gated populations is displayed in Figure 1J**-J’’**. **(I)** Representative LG expressing Fly-FUCCI in dome-positive progenitors (*dome>FUCCI*) with overexpression of Cdc25 (*UAS-Cdc25*), which results in a large increase in the number of G1 (green) cells as compared to *WT* (LG shown in Figure 1E). **(J-K)** Quantitative analysis of cell cycle phases G2 **(J)** and G1 **(K)** in individual progenitors (*dome>FUCCI*). Compared to *WT*, overexpression of Cdc25 (*UAS-Cdc25*) significantly decreases the number of progenitors in G2 **(J)** and significantly increases the number of progenitors in G1 **(K)**.

**Figure S2:**
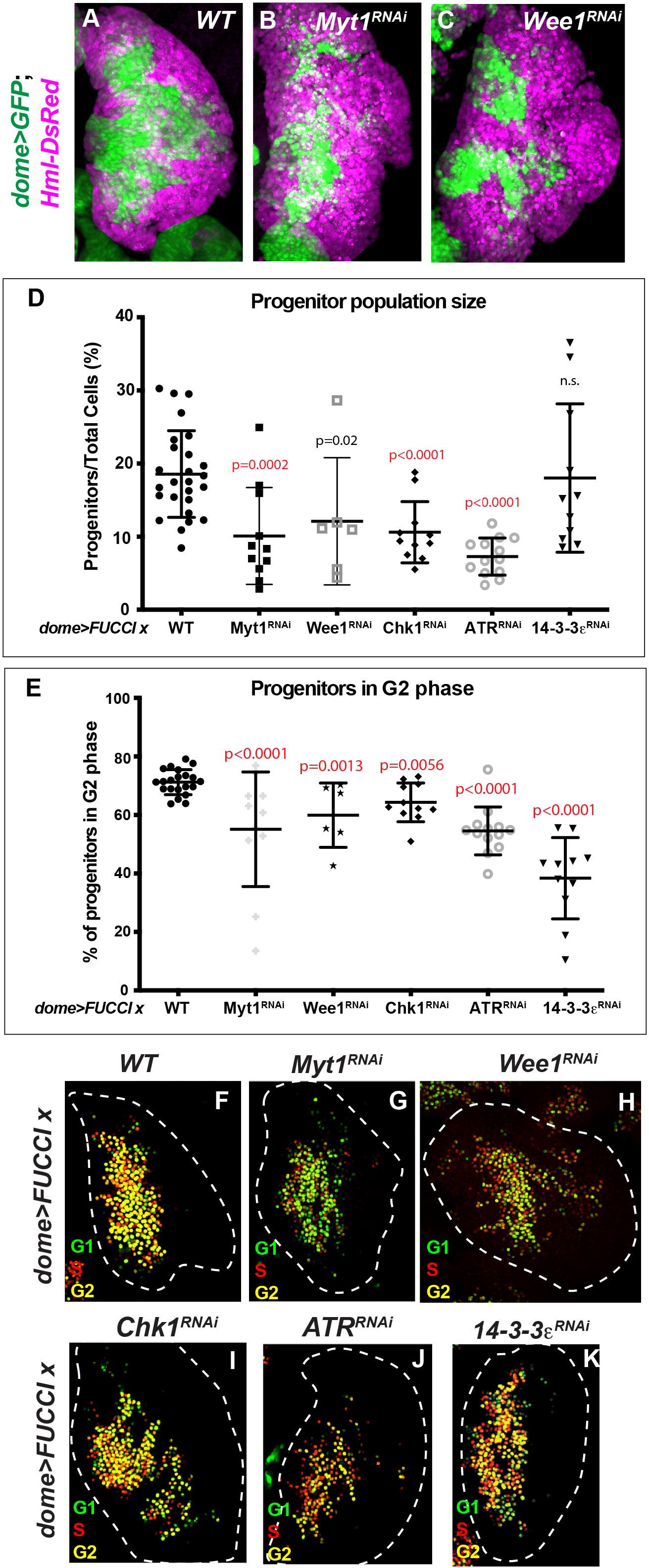
Prolonged G2 phase is important for maintaining the progenitor population and this pathway involves both classic and DNA damage-related cell cycle regulators. **(A-B)** Compared to wild type (WT) **(A)** (same image as in Figure 2A), RNAi-mediated depletion of classic G2/M regulators Myt1 (**B**) or Wee1 (**C**), in the progenitors (*dome>GFP*; green) decreases the progenitor population and increases the differentiated cells (*Hml-DsRed*; magenta). Middle third z-stack projections of representative lymph glands (LGs). **(D)** Quantitative analysis of the progenitor population from *dome>FUCCI* LGs. Compared to WT, RNAi-mediated loss of Myt1, Wee1, Chk1, and ATR all significantly decrease the size of the progenitor population. **(E)** Quantitative analysis of the cell cycle profile of progenitors (*dome>FUCCI*). Compared to WT, RNAi-mediated loss of Myt1, Wee1, Chk1, ATR, and 14-3-3ε all significantly decrease the fraction or percentage of progenitors in G2 phase. **(F-K)** Representative LGs expressing Fly-FUCCI in *dome*-positive progenitors (*dome>FUCCI*) from the genotypes analyzed in **(D-E)**.

**Figure S3:**
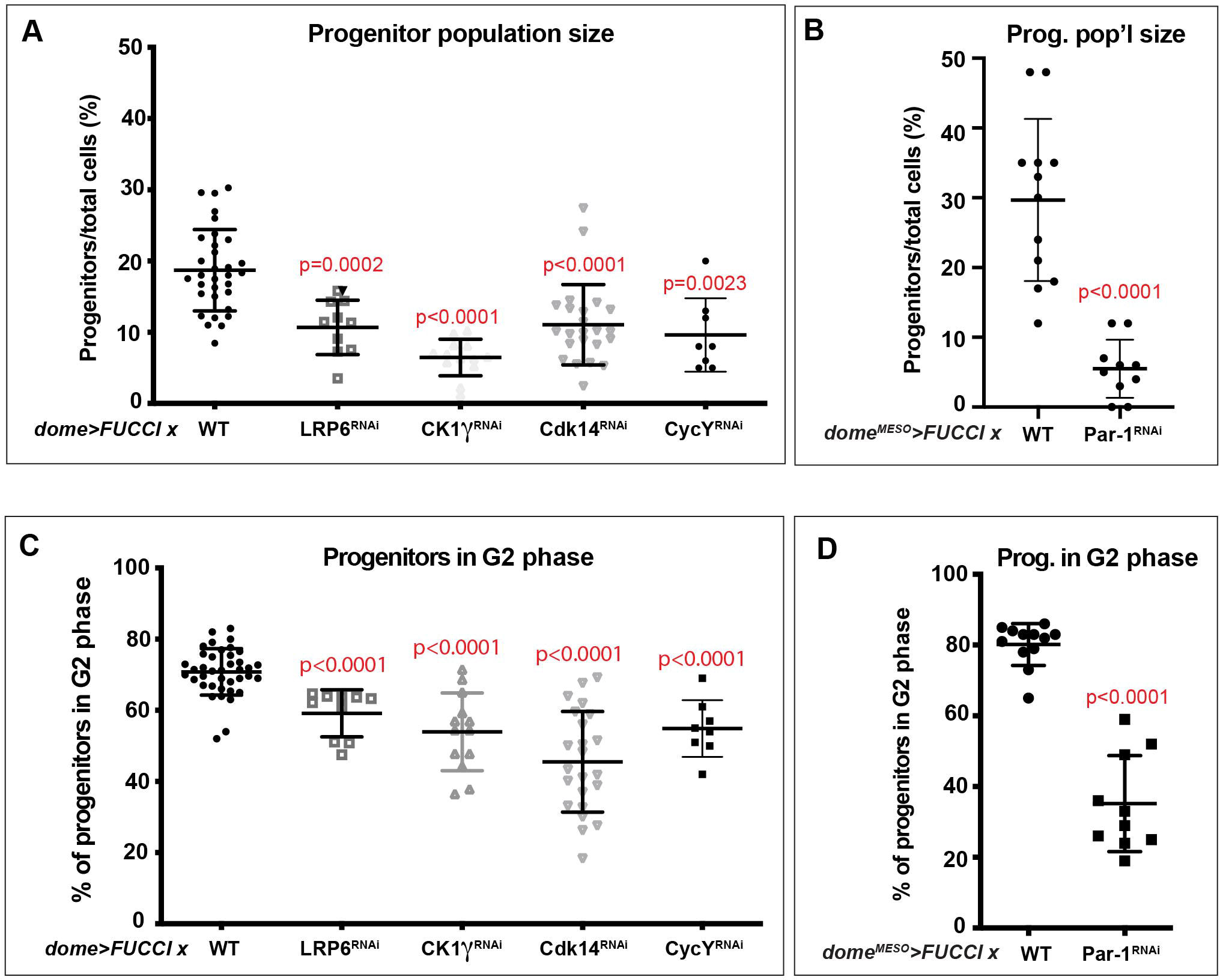
Wnt signaling pathway components function to control progenitor maintenance and G2 cell cycle progression. **(A)** Quantitative analysis of progenitor population size from *dome>FUCCI* LGs. Compared to wild type (WT), RNAi-mediated loss of Wnt signaling pathway components Arrow/LRP6, CK1γ, CDK14/Eip63E, and Cyclin Y all significantly decrease the size of the progenitor population. **(B)** Quantitative analysis of progenitor population size from *dome^MESO^>FUCCI* LGs. Compared to WT, *Par-1-RNAi* significantly decreases the size of the progenitor population. **(C)** Quantitative analysis of the cell cycle profile of progenitors from *dome>FUCCI* LGs. Compared to WT, RNAi-mediated loss of Wnt signaling pathway components Arrow/LRP6, CK1γ, CDK14/Eip63E, and Cyclin Y all significantly decrease the percentage of progenitors in G2 phase. **(D)** Quantitative analysis of the cell cycle profile of progenitors from *dome^MESO^>FUCCI* LGs. Compared to WT, *Par-1-RNAi* significantly decrease the percentage of progenitors in G2 phase.

**Figure S4:**
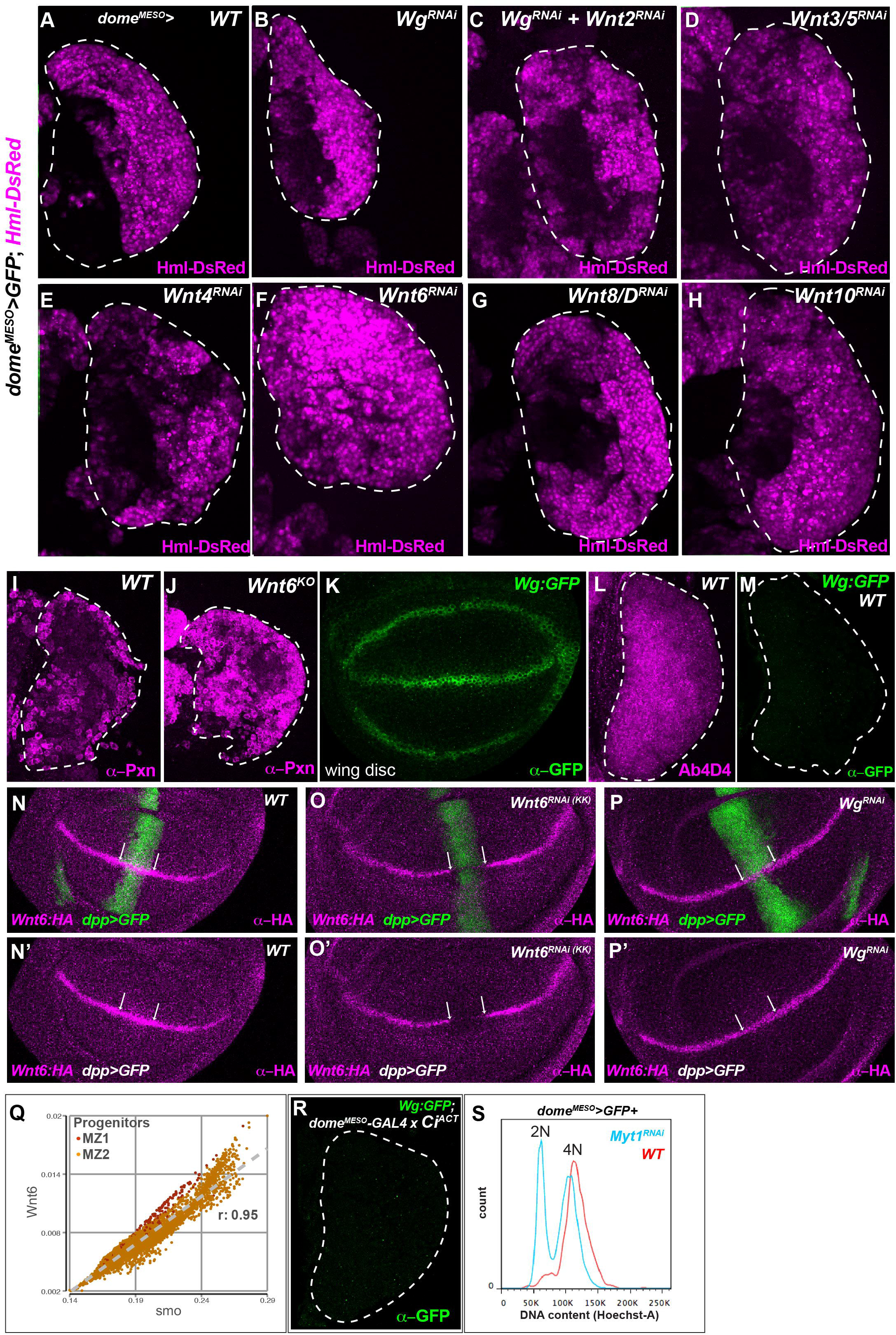
Wnt6 maintains progenitors in G2. **(A-H)** Compared to wild type **(A)**, RNAi-mediated depletion of Wnt ligands Wg **(B)**, Wg and Wnt2 **(C)**, Wnt3/5 **(D)**, Wnt4 **(E)**, Wnt8/D **(G)**, or Wnt10 **(H)** in the progenitors (*dome^MESO^-GAL4*) does not increase differentiation (*Hml-DsRed*; magenta). Only the loss of Wnt6 **(F)** increases differentiation. **(I-J)** Compared with wild type (*w*^1118^) **(I)**, loss of Wnt6 (knockout line *Wnt6^KO^*) **(J)** results in increased differentiation as indicated by anti-Pxn staining to mark differentiating cells. **(K)** Immunostaining with anti-GFP shows Wg:GFP expression along the D/V boundary of the wing imaginal disc. **(L)** Ab4D4 (anti-Wnt) shows high immunostaining in the progenitor population in wild-type LGs. **(M)** Immunostaining with anti-GFP shows Wg:GFP expression is not detected in wild-type lymph glands from 3rd instar larvae. Genotype: *Wg:GFP; dome^MESO^-GAL4*. **(N-P’)** Immunostaining with anti-HA shows Wnt6:HA is expressed along the D/V boundary in a wild-type wing disc **(N, N’)**. Expression of *UAS-Wnt6-RNAi* **(O, O’)** with *dpp>GFP* expressed along the A/P boundary results in loss of expression of Wnt6:HA within the intersecting region (white arrows). Wnt6:HA expression is not altered when *UAS-Wg-RNAi* **(P, P’)** is expressed along the A/P boundary. **(Q)** Wnt6 and smo transcript levels within individual progenitor cells show high correlation (r=0.95). Wnt6 transcript levels (normalized counts on the y-axis) and smo transcript levels (normalized counts on the x-axis) in the two progenitor populations identified in Girard *et al*.^41^ (single cell RNA sequencing results; MZ1 and MZ2). **(R)** Compared to WT **(M)**, overactivation of Hh signaling with a constitutively active Ci (*dome^MESO^-GAL4*; *UAS-Ci^ACT^*), does not change Wg:GFP levels in the lymph gland. **(S)** Flow cytometric DNA content analysis. Compared to WT (red), RNAi-mediated loss of Myt1 (blue) in the progenitors (*dome^MESO^>GFP*) drastically changes the cell cycle profile of progenitors from predominantly in G2 (4N) in WT to largely in G1 (2N) in Myt1-RNAi LGs.

**Figure S5:**
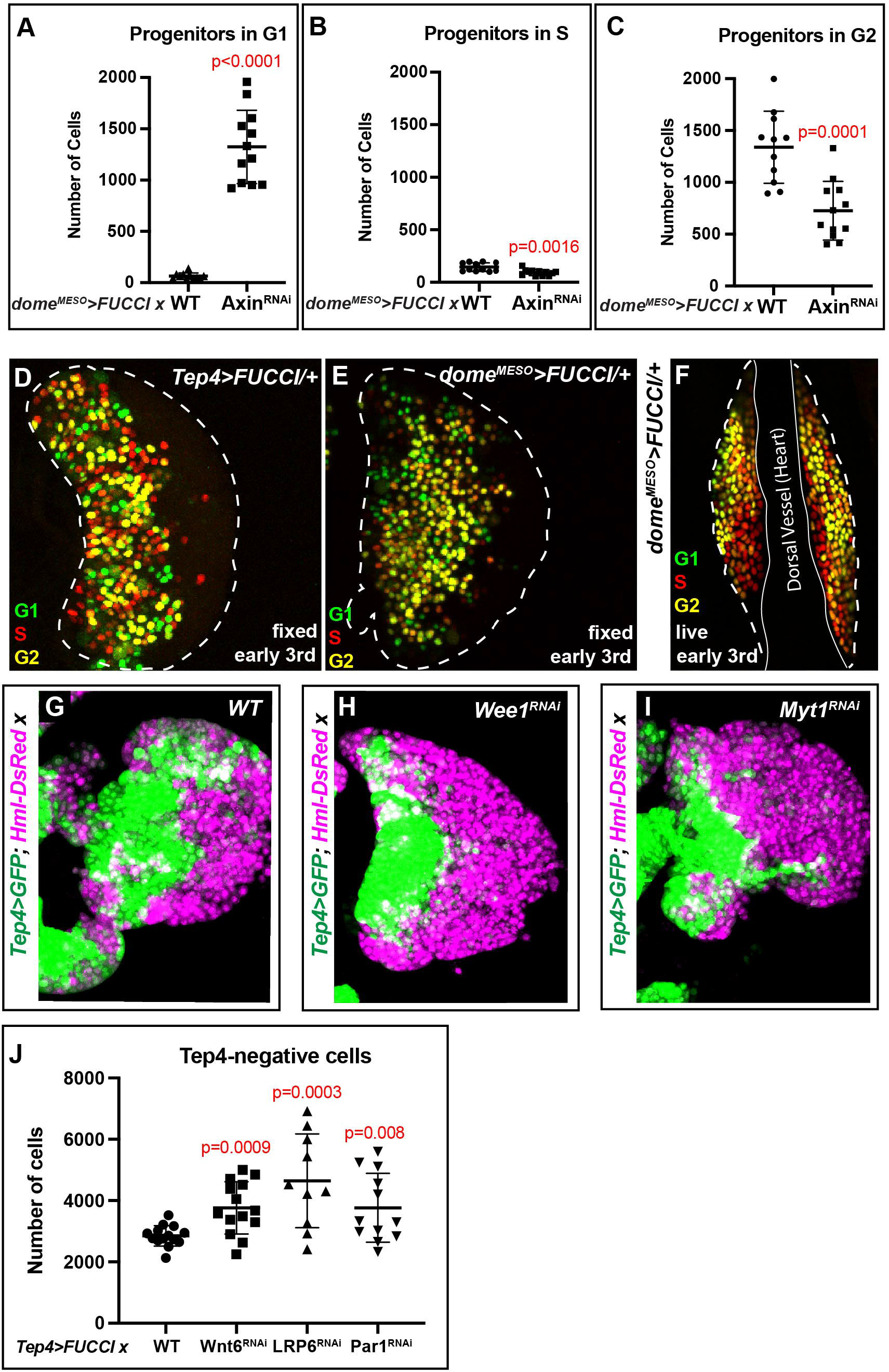
Role of β-catenin in cell cycle regulation. **(A-C)** Quantitation of cell cycle data (*dome^MESO^>FUCCI*) demonstrates that compared to wild type, RNAi-mediated loss of Axin (Axin^RNAi^) (activation of the Wnt pathway) results in a significant increase in the number of progenitors in G1 phase **(A)**, a significant decrease in the number of progenitors in S phase **(B)**, and a significant decrease in the number of progenitors in G2 phase **(C)**. **(D-E)** Representative images of fixed wild-type lymph glands from early 3rd instar expressing *Tep4-GAL4, UAS-FUCCI* **(D)** or *dome^MESO^-GAL4, UAS-FUCCI* **(E)**. **(F)** Live imaging of a lymph gland from an early 3rd instar larva expressing *dome^MESO^>FUCCI* shows that while most progenitors are in G2 (yellow), core progenitors near the heart (outlined with a solid white line) can be seen in S phase (red) in early 3rd instar larvae. **(G-I)** Compared to WT **(G)**, RNAi–mediated loss of Wee1 **(H)** or Myt1 **(I)** in *Tep4*-positive progenitors (*Tep4>GFP*) increases differentiated cells (*Hml-DsRed;* magenta) and decreases *Tep4*-positive progenitors (green). **(J)** Quantitation of the number of the differentiating cells (*Tep4*-negative cells) in lymph glands expressing *Tep4>FUCCI*. Compared with wild type (WT), RNAi–mediated loss of Wnt6, LRP6, or Par-1 significantly increases the number of differentiating cells.

**Figure S6:**
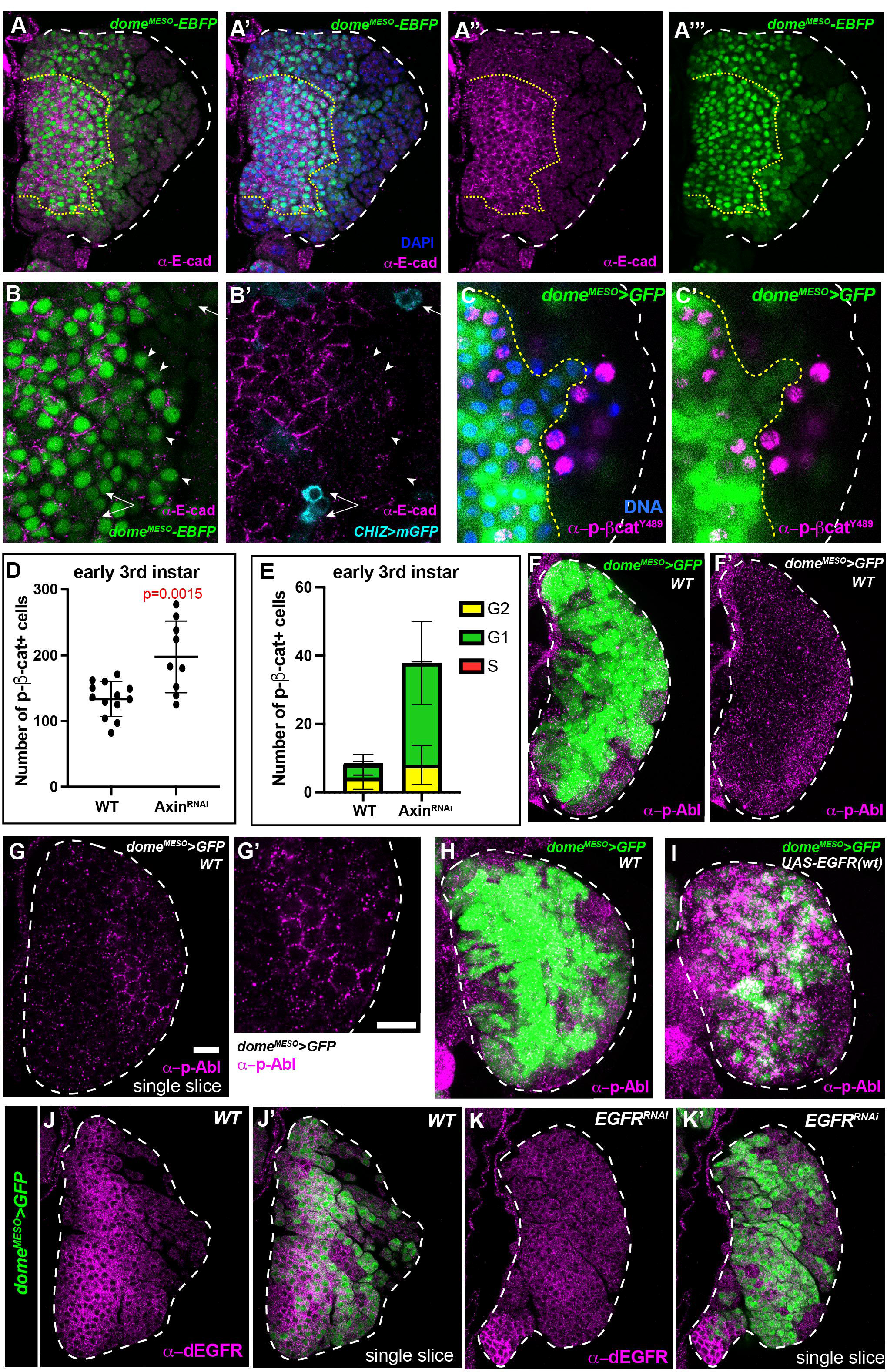
E-Cadherin and dual function of β-catenin in spatially restricted zones. **(A-A’’’)** E-cadherin-negative, *dome^MESO^-EBFP* expressing progenitors (green) can be seen outside of the high E-cadherin (magenta) expression region (yellow dotted line) nearer to the distal edge of the lymph gland (white dashed line). **(B,B’)** Higher magnification views show that most E-cadherin-negative, *dome^MESO^*-positive progenitors (arrowheads) are not intermediate progenitors as they lack *CHIZ-GAL4, UAS-mGFP* (cyan; arrows), a marker of the intermediate zone^42^. Immunostaining with anti-E-cadherin antibody (magenta). **(C,C’)** Immunostaining of wild-type lymph glands expressing *dome^MESO^-GAL4, UAS-GFP* (green) with anti-pY489-β-cat (magenta) shows that phospho-β-catenin is localized to the nucleus of some progenitors (green) closer to the distal edge of the progenitor population (yellow dotted line) as well as GFP-negative differentiating cells (which no longer express *dome^MESO^-GAL4*) near the distal edge of the lymph gland (white dashed line). DAPI/DNA (blue). **(D)** Quantitation of data from Figures 6B**,B’** and **6G,G’** shows a significant increase in the total number of pY489-β-catenin-positive cells upon RNAi-mediated loss of Axin. **(E)** Quantitation of FUCCI results (*dome^MESO^>FUCCI* immunostained with anti-pY489-β-cat) shows that, compared to WT, a significant increase in the number of pY489-β-cat-positive progenitors in G1 is seen upon RNAi-mediated loss of Axin with a larger fraction of pY489-β-cat-positive progenitors in G1 than in G2. **(F-F’)** pY412-Abl levels increase in progenitors (*dome^MESO^>GFP*; green) and differentiating cells (*dome^MESO^*-negative) near the distal edge of the lymph gland (white dotted line). Maximum intensity projection of several confocal slices. **(G-G’)** single confocal slice shows anti-pY412-Abl staining (magenta) in puncta in the cytoplasm and in a honeycomb pattern at the cell surface. **(G’)** higher magnification view. **(H,I)** Corresponding images from Figure 6M**,N** displaying fluorescent channels for both *dome^MESO^>GFP* (green) and anti-pY412-Abl staining (magenta). **(J-K’)** In wild-type lymph glands **(J-J’)** high levels of dEGFR staining (*Drosophila* EGFR; magenta) can be seen throughout the progenitor population (*dome^MESO^>GFP*; green) but is largely reduced upon RNAi-mediated loss of EGFR **(K-K’)**. Single confocal slices.

## References

1. Hartwell, L.H., and Weinert, T.A. (1989). Checkpoints: controls that ensure the order of cell cycle events. Science 246, 629–634. 10.1126/science.2683079.

2. Hochegger, H., Takeda, S., and Hunt, T. (2008). Cyclin-dependent kinases and cell-cycle transitions: does one fit all? Nat. Rev. Mol. Cell Biol. 9, 910–916. 10.1038/nrm2510.

3. Basu, S., Greenwood, J., Jones, A.W., and Nurse, P. (2022). Core control principles of the eukaryotic cell cycle. Nature 607, 381–386. 10.1038/s41586-022-04798-8.

4. Lee, L.A., and Orr-Weaver, T.L. (2003). Regulation of Cell Cycles in Drosophila Development: Intrinsic and Extrinsic Cues. Annu. Rev. Genet. 37, 545–578. 10.1146/annurev.genet.37.110801.143149.

5. Kimata, Y., Leturcq, M., and Aradhya, R. (2020). Emerging roles of metazoan cell cycle regulators as coordinators of the cell cycle and differentiation. FEBS Lett. 10.1002/1873-3468.13805.

6. Marescal, O., and Cheeseman, I.M. (2020). Cellular Mechanisms and Regulation of Quiescence. Dev. Cell 55, 259–271. 10.1016/j.devcel.2020.09.029.

7. Spencer, S.L., Cappell, S.D., Tsai, F.-C., Overton, K.W., Wang, C.L., and Meyer, T. (2013). The proliferation-quiescence decision is controlled by a bifurcation in CDK2 activity at mitotic exit. Cell 155, 369–383. 10.1016/j.cell.2013.08.062.

8. Cheung, T.H., and Rando, T.A. (2013). Molecular regulation of stem cell quiescence. Nat. Rev. Mol. Cell Biol. 14, 329–340. 10.1038/nrm3591.

9. Pauklin, S., and Vallier, L. (2013). The cell-cycle state of stem cells determines cell fate propensity. Cell 155, 135–147. 10.1016/j.cell.2013.08.031.

10. Jang, J., Han, D., Golkaram, M., Audouard, M., Liu, G., Bridges, D., Hellander, S., Chialastri, A., Dey, S.S., Petzold, L.R., et al. (2019). Control over single-cell distribution of G1 lengths by WNT governs pluripotency. PLOS Biol. 17, e3000453. 10.1371/journal.pbio.3000453.

11. Lange, C., and Calegari, F. (2010). Cdks and cyclins link G1 length and differentiation of embryonic, neural and hematopoietic stem cells. Cell Cycle 9, 1893–1900. 10.4161/cc.9.10.11598.

12. Soufi, A., and Dalton, S. (2016). Cycling through developmental decisions: how cell cycle dynamics control pluripotency, differentiation and reprogramming. Dev. Camb. Engl. 143, 4301–4311. 10.1242/dev.142075.

13. Marechal, A., and Zou, L. (2013). DNA Damage Sensing by the ATM and ATR Kinases. Cold Spring Harb. Perspect. Biol. 5, a012716–a012716. 10.1101/cshperspect.a012716.

14. Smits, V.A.J., and Gillespie, D.A. (2015). DNA damage control: regulation and functions of checkpoint kinase 1. FEBS J. 282, 3681–3692. 10.1186/gb-2011-12-8-r78.

15. Egger, B., Gold, K.S., and Brand, A.H. (2011). Regulating the balance between symmetric and asymmetric stem cell division in the developing brain. Fly (Austin) 5, 237–241. 10.4161/fly.5.3.15640.

16. Venkei, Z.G., and Yamashita, Y.M. (2018). Emerging mechanisms of asymmetric stem cell division. J. Cell Biol. 217, 3785–3795. 10.1083/jcb.201807037.

17. Rim, E.Y., Clevers, H., and Nusse, R. (2022). The Wnt Pathway: From Signaling Mechanisms to Synthetic Modulators. Annu. Rev. Biochem. 91, 571–598. 10.1146/annurev-biochem-040320-103615.

18. Acebron, S.P., Karaulanov, E., Berger, B.S., Huang, Y.-L., and Niehrs, C. (2014). Mitotic wnt signaling promotes protein stabilization and regulates cell size. Mol. Cell 54, 663–674. 10.1016/j.molcel.2014.04.014.

19. Davidson, G., Shen, J., Huang, Y.-L., Su, Y., Karaulanov, E., Bartscherer, K., Hassler, C., Stannek, P., Boutros, M., and Niehrs, C. (2009). Cell Cycle Control of Wnt Receptor Activation. Dev. Cell 17, 788–799. 10.1016/j.devcel.2009.11.006.

20. Davidson, G., and Niehrs, C. (2010). Emerging links between CDK cell cycle regulators and Wnt signaling. Trends Cell Biol. 20, 453–460. 10.1016/j.tcb.2010.05.002.

21. Huang, Y.-L., Anvarian, Z., Döderlein, G., Acebron, S.P., and Niehrs, C. (2015). Maternal Wnt/STOP signaling promotes cell division during early Xenopusembryogenesis. Proc. Natl. Acad. Sci. U. S. A. 112, 5732–5737. 10.1038/349132a0.

22. Valenta, T., Hausmann, G., and Basler, K. (2012). The many faces and functions of β-catenin. EMBO J. 31, 2714–2736. 10.1038/emboj.2012.150.

23. Tepass, U., Gruszynski-DeFeo, E., Haag, T.A., Omatyar, L., Török, T., and Hartenstein, V. (1996). shotgun encodes Drosophila E-cadherin and is preferentially required during cell rearrangement in the neurectoderm and other morphogenetically active epithelia. Genes Dev. 10, 672–685. 10.1101/gad.10.6.672.

24. Meng, W., and Takeichi, M. (2009). Adherens Junction: Molecular Architecture and Regulation. Cold Spring Harb. Perspect. Biol. 1, a002899. 10.1101/cshperspect.a002899.

25. Heuberger, J., and Birchmeier, W. (2010). Interplay of cadherin-mediated cell adhesion and canonical Wnt signaling. Cold Spring Harb. Perspect. Biol. 2, a002915. 10.1101/cshperspect.a002915.

26. Evans, C.J., Hartenstein, V., and Banerjee, U. (2003). Thicker than blood: conserved mechanisms in Drosophila and vertebrate hematopoiesis. Dev. Cell 5, 673–690.

27. Hartenstein, V. (2006). Blood Cells and Blood Cell Development in the Animal Kingdom. Annu. Rev. Cell Dev. Biol. 22, 677–712. 10.1146/annurev.cellbio.22.010605.093317.

28. Grigorian, M., Mandal, L., and Hartenstein, V. (2011). Hematopoiesis at the onset of metamorphosis: terminal differentiation and dissociation of the Drosophila lymph gland. Dev. Genes Evol. 221, 121–131. 10.1007/s00427-011-0364-6.

29. Lebestky, T., Chang, T., Hartenstein, V., and Banerjee, U.B. (2000). Specification of Drosophila hematopoietic lineage by conserved transcription factors. Science 288, 146–149. 10.1126/science.288.5463.146.

30. Banerjee, U., Girard, J.R., Goins, L.M., and Spratford, C.M. (2019). Drosophila as a Genetic Model for Hematopoiesis. Genetics 211, 367–417. 10.1534/genetics.118.300223.

31. Crozatier, M., Ubeda, J.-M., Vincent, A., and Meister, M. (2004). Cellular immune response to parasitization in Drosophila requires the EBF orthologue collier. PLoS Biol. 2, E196. 10.1371/journal.pbio.0020196.

32. Jung, S.-H., Evans, C.J., Uemura, C., and Banerjee, U. (2005). The Drosophila lymph gland as a developmental model of hematopoiesis. Dev. Camb. Engl. 132, 2521–2533. 10.1242/dev.01837.

33. Mandal, L., Martinez-Agosto, J.A., Evans, C.J., Hartenstein, V., and Banerjee, U. (2007). A Hedgehog-and Antennapedia-dependent niche maintains Drosophila haematopoietic precursors. Nature 446, 320–324. 10.1038/nature05585.

34. Morin-Poulard, I. Euml l, Sharma, A., Louradour, I., Vanzo, N., Vincent, A., and le Crozatier, M. egrave (2016). Vascular control of the Drosophila haematopoietic microenvironment by Slit/Robo signalling. Nat. Commun. 7, 1–12. 10.1038/ncomms11634.

35. Destalminil-Letourneau, M., Morin-Poulard, I., Tian, Y., Vanzo, N., and Crozatier, M. (2021). The vascular niche controls Drosophila hematopoiesis via fibroblast growth factor signaling. eLife 10, e64672. 10.7554/eLife.64672.

36. Benmimoun, B., Polesello, C., Haenlin, M., and Waltzer, L. (2015). The EBF transcription factor Collier directly promotes Drosophila blood cell progenitor maintenance independently of the niche. Proc. Natl. Acad. Sci. U. S. A. 112, 9052–9057. 10.1073/pnas.1423967112.

37. Oyallon, J., Vanzo, N., Krzemien, J., Morin-Poulard, I., Vincent, A., and Crozatier, M. (2016). Two Independent Functions of Collier/Early B Cell Factor in the Control of Drosophila Blood Cell Homeostasis. PLOS ONE 11, e0148978. 10.1371/journal.pone.0148978.

38. Baldeosingh, R., Gao, H., Wu, X., and Fossett, N. (2018). Hedgehog signaling from the Posterior Signaling Center maintains U-shaped expression and a prohemocyte population in Drosophila. Dev. Biol. 441, 132–145. 10.1016/j.ydbio.2018.06.020.

39. Blanco-Obregon, D., Katz, M.J., Durrieu, L., Gándara, L., and Wappner, P. (2020). Context-specific functions of Notch in Drosophila blood cell progenitors. Dev. Biol. 462, 101–115. 10.1016/j.ydbio.2020.03.018.

40. Cho, B., Yoon, S.-H., Lee, D., Koranteng, F., Tattikota, S.G., Cha, N., Shin, M., Do, H., Hu, Y., Oh, S.Y., et al. (2020). Single-cell transcriptome maps of myeloid blood cell lineages in Drosophila. Nat. Commun., 1–18. 10.1038/s41467-020-18135-y.

41. Girard, J.R., Goins, L.M., Vuu, D.M., Sharpley, M.S., Spratford, C.M., Mantri, S.R., and Banerjee, U. (2021). Paths and pathways that generate cell-type heterogeneity and developmental progression in hematopoiesis. eLife 10, e67516. 10.7554/eLife.67516.

42. Spratford, C.M., Goins, L.M., Chi, F., Girard, J.R., Macias, S.N., Ho, V.W., and Banerjee, U. (2021). Intermediate progenitor cells provide a transition between hematopoietic progenitors and their differentiated descendants. Development 148, dev200216. 10.1242/dev.200216.

43. Mondal, B.C., Mukherjee, T., Mandal, L., Evans, C.J., Sinenko, S.A., Martinez-Agosto, J.A., and Banerjee, U. (2011). Interaction between differentiating cell-and niche-derived signals in hematopoietic progenitor maintenance. Cell 147, 1589–1600. 10.1016/j.cell.2011.11.041.

44. Letourneau, M., Lapraz, F., Sharma, A., Vanzo, N., Waltzer, L., and Crozatier, M. (2016). Drosophilahematopoiesis under normal conditions and in response to immune stress. FEBS Lett. 590, 4034–4051. 10.1182/blood-2014-05-577940.

45. Yu, S., Luo, F., and Jin, L.H. (2018). The Drosophila lymph gland is an ideal model for studying hematopoiesis. Dev. Comp. Immunol. 83, 60–69. 10.1016/j.dci.2017.11.017.

46. Sharma, S.K., Ghosh, S., Geetha, A.R., Mandal, S., and Mandal, L. (2019). Cell Adhesion-Mediated Actomyosin Assembly Regulates the Activity of Cubitus Interruptus for Hematopoietic Progenitor Maintenance in Drosophila. Genetics 212, 1279–1300. 10.1534/genetics.119.302209.

47. Kapoor, A., Padmavathi, A., Madhwal, S., and Mukherjee, T. (2022). Dual control of dopamine in Drosophila myeloid-like progenitor cell proliferation and regulation of lymph gland growth. EMBO Rep. 23, e52951. 10.15252/embr.202152951.

48. Zielke, N., Korzelius, J., van Straaten, M., Bender, K., Schuhknecht, G.F.P., Dutta, D., Xiang, J., and Edgar, B.A. (2014). Fly-FUCCI: A Versatile Tool for Studying Cell Proliferation in Complex Tissues. CellReports 7, 588–598. 10.1016/j.celrep.2014.03.020.

49. Lehman, D.A., Patterson, B., Johnston, L.A., Balzer, T., Britton, J.S., Saint, R., and Edgar, B.A. (1999). Cis-regulatory elements of the mitotic regulator, string/Cdc25. Dev. Camb. Engl. 126, 1793–1803.

50. Edgar, B.A., and O’Farrell, P.H. (1989). Genetic control of cell division patterns in the Drosophila embryo. Cell 57, 177–187. 10.1016/0092-8674(89)90183-9.

51. Kimelman, D. (2014). Cdc25 and the importance of G2 control. Cell Cycle 13, 2165–2171. 10.4161/cc.29537.

52. Perry, J.A., and Kornbluth, S. (2007). Cdc25 and Wee1: analogous opposites? Cell Div. 2, 12. 10.1186/1747-1028-2-12.

53. Hutchins, J.R.A., and Clarke, P.R. (2004). Many fingers on the mitotic trigger: post-translational regulation of the Cdc25C phosphatase. Cell Cycle 3, 41–45.

54. Wells, N.J., Watanabe, N., Tokusumi, T., Jiang, W., Verdecia, M.A., and Hunter, T. (1999). The C-terminal domain of the Cdc2 inhibitory kinase Myt1 interacts with Cdc2 complexes and is required for inhibition of G(2)/M progression. J. Cell Sci. 112 (Pt 19), 3361–3371.

55. Niida, H., and Nakanishi, M. (2006). DNA damage checkpoints in mammals. Mutagenesis 21, 3–9. 10.1093/mutage/gei063.

56. Abraham, R.T. (2001). Cell cycle checkpoint signaling through the ATM and ATR kinases. Genes Dev. 15, 2177–2196. 10.1101/gad.914401.

57. Zhao, H., and Piwnica-Worms, H. (2001). ATR-Mediated Checkpoint Pathways Regulate Phosphorylation and Activation of Human Chk1. Mol. Cell. Biol. 21, 4129–4139. 10.1128/MCB.21.13.4129-4139.2001.

58. Vries, H.I. de, Uyetake, L., Lemstra, W., Brunsting, J.F., Su, T.T., Kampinga, H.H., and Sibon, O.C.M. (2005). Grp/DChk1 is required for G2-M checkpoint activation in Drosophila S2 cells, whereas Dmnk/DChk2 is dispensable. J. Cell Sci. 118, 1833–1842. 10.1242/jcs.02309.

59. Liu, Q., Guntuku, S., Cui, X.S., Matsuoka, S., Cortez, D., Tamai, K., Luo, G., Carattini-Rivera, S., DeMayo, F., Bradley, A., et al. (2000). Chk1 is an essential kinase that is regulated by Atr and required for the G(2)/M DNA damage checkpoint. Genes Dev. 14, 1448–1459.

60. Zachos, G., and Gillespie, D. (2007). Exercising Restraints: Role of Chk1 in Regulating the Onset and Progression of Unperturbed Mitosis in Vertebrate Cells. Cell Cycle 6, 810–813. 10.4161/cc.6.7.4048.

61. Su, T.T., Parry, D.H., Donahoe, B., Chien, C.T., O’Farrell, P.H., and Purdy, A. (2001). Cell cycle roles for two 14-3-3 proteins during Drosophila development. J. Cell Sci. 114, 3445–3454.

62. Lee, J., Kumagai, A., and Dunphy, W.G. (2001). Positive regulation of Wee1 by Chk1 and 14-3-3 proteins. Mol. Biol. Cell 12, 551–563.

63. Sørensen, C.S., and Syljuåsen, R.G. (2012). Safeguarding genome integrity: the checkpoint kinases ATR, CHK1 and WEE1 restrain CDK activity during normal DNA replication. Nucleic Acids Res. 40, 477–486. 10.1093/nar/gkr697.

64. Brunet, A., Kanai, F., Stehn, J., Xu, J., Sarbassova, D., Frangioni, J.V., Dalal, S.N., DeCaprio, J.A., Greenberg, M.E., and Yaffe, M.B. (2002). 14-3-3 transits to the nucleus and participates in dynamic nucleocytoplasmic transport. J. Cell Biol. 156, 817–828. 10.1083/jcb.200112059.

65. Göransson, O., Deak, M., Wullschleger, S., Morrice, N.A., Prescott, A.R., and Alessi, D.R. (2006). Regulation of the polarity kinases PAR-1/MARK by 14-3-3 interaction and phosphorylation. J. Cell Sci. 119, 4059–4070. 10.1242/jcs.03097.

66. Benton, R., and St Johnston, D. (2003). Drosophila PAR-1 and 14-3-3 inhibit Bazooka/PAR-3 to establish complementary cortical domains in polarized cells. Cell 115, 691–704. 10.1016/s0092-8674(03)00938-3.

67. Benton, R., Palacios, I.M., and St Johnston, D. (2002). Drosophila 14-3-3/PAR-5 is an essential mediator of PAR-1 function in axis formation. Dev. Cell 3, 659–671. 10.1016/S1534-5807(02)00320-9.

68. Sun, T.Q., Lu, B., Feng, J.J., Reinhard, C., Jan, Y.N., Fantl, W.J., and Williams, L.T. (2001). PAR-1 is a Dishevelled-associated kinase and a positive regulator of Wnt signalling. Nat. Cell Biol. 3, 628–636. 10.1038/35083016.

69. Ossipova, O., Dhawan, S., Sokol, S., and Green, J.B.A. (2005). Distinct PAR-1 proteins function in different branches of Wnt signaling during vertebrate development. Dev. Cell 8, 829–841. 10.1016/j.devcel.2005.04.011.

70. Tolwinski, N.S., Wehrli, M., Rives, A., Erdeniz, N., DiNardo, S., and Wieschaus, E. (2003). Wg/Wnt signal can be transmitted through arrow/LRP5,6 and Axin independently of Zw3/Gsk3beta activity. Dev. Cell 4, 407–418.

71. Davidson, G., Wu, W., Shen, J., Bilic, J., Fenger, U., Stannek, P., Glinka, A., and Niehrs, C. (2005). Casein kinase 1 gamma couples Wnt receptor activation to cytoplasmic signal transduction. Nature 438, 867–872. 10.1038/nature04170.

72. Jiang, J. (2017). CK1 in Developmental signaling: Hedgehog and Wnt. Curr. Top. Dev. Biol. 123, 303–329. 10.1016/bs.ctdb.2016.09.002.

73. Liu, D., and Finley, R.L., Jr. (2010). Cyclin Y Is a Novel Conserved Cyclin Essential for Development in Drosophila. Genetics 184, 1025–1035. 10.1534/genetics.110.114017.

74. Liu, D., Guest, S., and Finley, R.L. (2010). Why Cyclin Y? A highly conserved cyclin with essential functions. Fly (Austin) 4, 278–282. 10.4161/fly.4.4.12881.

75. Li, S., Jiang, M., Wang, W., and Chen, J. (2014). 14-3-3 Binding to Cyclin Y contributes to cyclin Y/CDK14 association. Acta Biochim. Biophys. Sin. 46, 299–304. 10.1093/abbs/gmu005.

76. Swarup, S., and Verheyen, E.M. (2012). Wnt/Wingless signaling in Drosophila. Cold Spring Harb. Perspect. Biol. 4. 10.1101/cshperspect.a007930.

77. Peng, C.Y., Graves, P.R., Ogg, S., Thoma, R.S., Byrnes, M.J., Wu, Z., Stephenson, M.T., and Piwnica-Worms, H. (1998). C-TAK1 protein kinase phosphorylates human Cdc25C on serine 216 and promotes 14-3-3 protein binding. Cell Growth Amp Differ. Mol. Biol. J. Am. Assoc. Cancer Res. 9, 197–208.

78. Müller, J., Ritt, D.A., Copeland, T.D., and Morrison, D.K. (2003). Functional analysis of C-TAK1 substrate binding and identification of PKP2 as a new C-TAK1 substrate. EMBO J. 22, 4431–4442. 10.1093/emboj/cdg426.

79. Kizhedathu, A., Kunnappallil, R.S., Bagul, A.V., Verma, P., and Guha, A. (2020). Multiple Wnts act synergistically to induce Chk1/Grapes expression and mediate G2 arrest in Drosophila tracheoblasts. eLife 9, e57056. 10.7554/eLife.57056.

80. Brook, W.J., and Cohen, S.M. (1996). Antagonistic Interactions Between Wingless and Decapentaplegic Responsible for Dorsal-Ventral Pattern in the Drosophila Leg. Science 273, 1373–1377. 10.1126/science.273.5280.1373.

81. Shim, J., Mukherjee, T., and Banerjee, U. (2012). Direct sensing of systemic and nutritional signals by haematopoietic progenitors in Drosophila. Nat. Cell Biol. 14, 394–400. 10.1038/ncb2453.

82. Sinenko, S.A., Mandal, L., Martinez-Agosto, J.A., and Banerjee, U. (2009). Dual role of wingless signaling in stem-like hematopoietic precursor maintenance in Drosophila. Dev. Cell 16, 756–763. 10.1016/j.devcel.2009.03.003.

83. Zhang, C.U., Blauwkamp, T.A., Burby, P.E., and Cadigan, K.M. (2014). Wnt-mediated repression via bipartite DNA recognition by TCF in the Drosophila hematopoietic system. PLOS Genet. 10, e1004509. 10.1371/journal.pgen.1004509.

84. Doumpas, N., Jékely, G., and Teleman, A.A. (2013). Wnt6 is required for maxillary palp formation in Drosophila. BMC Biol. 11, 104. 10.1186/1741-7007-11-104.

85. Port, F., Chen, H.-M., Lee, T., and Bullock, S.L. (2014). Optimized CRISPR/Cas tools for efficient germline and somatic genome engineering in Drosophila. Proc. Natl. Acad. Sci. U. S. A. 111, E2967–76. 10.1073/pnas.1405500111.

86. Lee, K.-A., Kim, B., Bhin, J., Do Hun Kim, You, H., Kim, E.-K., Kim, S.-H., Ryu, J.-H., Hwang, D., and Lee, W.-J. (2015). Bacterial Uracil Modulates Drosophila DUOX-Dependent Gut Immunity via Hedgehog-Induced Signaling Endosomes. Cell Host Microbe 17, 191–204. 10.1016/j.chom.2014.12.012.

87. Tokusumi, Y., Tokusumi, T., Stoller-Conrad, J., and Schulz, R.A. (2010). Serpent, Suppressor of Hairless and U-shaped are crucial regulators of hedgehog niche expression and prohemocyte maintenance during Drosophila larval hematopoiesis. Development 137, 3561–3568. 10.1242/dev.053728.

88. Giordani, G., Barraco, M., Giangrande, A., Martinelli, G., Guadagnuolo, V., Simonetti, G., Perini, G., and Bernardoni, R. (2016). The human Smoothened inhibitor PF-04449913 induces exit from quiescence and loss of multipotent Drosophila hematopoietic progenitor cells. Oncotarget 7, 55313–55327. 10.18632/oncotarget.10879.

89. Tolwinski, N.S., and Wieschaus, E. (2001). Armadillo nuclear import is regulated by cytoplasmic anchor Axin and nuclear anchor dTCF/Pan. Development 128, 2107–2117. 10.1242/dev.128.11.2107.

90. Hamada, F., Tomoyasu, Y., Takatsu, Y., Nakamura, M., Nagai, S., Suzuki, A., Fujita, F., Shibuya, H., Toyoshima, K., Ueno, N., et al. (1999). Negative Regulation of Wingless Signaling by D-Axin, a Drosophila Homolog of Axin. Science 283, 1739–1742. 10.1126/science.283.5408.1739.

91. Willert, K., Shibamoto, S., and Nusse, R. (1999). Wnt-induced dephosphorylation of Axin releases β-catenin from the Axin complex. Genes Dev. 13, 1768–1773.

92. Krzemien, J., Dubois, L., Makki, R., Meister, M., Vincent, A., and Crozatier, M. (2007). Control of blood cell homeostasis in Drosophila larvae by the posterior signalling centre. Nature 446, 325–328. 10.1038/nature05650.

93. Irving, P., Ubeda, J.-M., Doucet, D., Troxler, L., Lagueux, M., Zachary, D., Hoffmann, J.A., Hetru, C., and Meister, M. (2005). New insights into Drosophila larval haemocyte functions through genome-wide analysis. Cell. Microbiol. 7, 335–350. 10.1111/j.1462-5822.2004.00462.x.

94. Hombría, J.C.-G., Brown, S., Häder, S., and Zeidler, M.P. (2005). Characterisation of Upd2, a Drosophila JAK/STAT pathway ligand. Dev. Biol. 288, 420–433. 10.1016/j.ydbio.2005.09.040.

95. Evans, C.J., Liu, T., and Banerjee, U. (2014). Drosophila hematopoiesis: Markers and methods for molecular genetic analysis. Methods San Diego Calif. 10.1016/j.ymeth.2014.02.038.

96. Gao, H., Wu, X., and Fossett, N. (2013). Drosophila E-cadherin functions in hematopoietic progenitors to maintain multipotency and block differentiation. PLOS ONE 8, e74684. 10.1371/journal.pone.0074684.

97. Gao, H., Wu, X., Simon, L., and Fossett, N. (2014). Antioxidants maintain E-cadherin levels to limit Drosophila prohemocyte differentiation. PLOS ONE 9, e107768. 10.1371/journal.pone.0107768.

98. van Roy, F., and Berx, G. (2008). The cell-cell adhesion molecule E-cadherin. Cell. Mol. Life Sci. 65, 3756–3788. 10.1007/s00018-008-8281-1.

99. Harris, T.J.C., and Tepass, U. (2010). Adherens junctions: from molecules to morphogenesis. Nat. Rev. Mol. Cell Biol. 11, 502–514. 10.1038/nrm2927.

100. Riggleman, B., Schedl, P., and Wieschaus, E. (1990). Spatial expression of the Drosophila segment polarity gene armadillo is posttranscriptionally regulated by wingless. Cell 63, 549–560. 10.1016/0092-8674(90)90451-J.

101. Lien, W.-H., and Fuchs, E. (2014). Wnt some lose some: transcriptional governance of stem cells by Wnt/β-catenin signaling. Genes Amp Dev. 28, 1517–1532. 10.1101/gad.244772.114.

102. Shah, K., and Kazi, J.U. (2022). Phosphorylation-Dependent Regulation of WNT/Beta-Catenin Signaling. Front. Oncol. 12.

103. Rhee, J., Buchan, T., Zukerberg, L., Lilien, J., and Balsamo, J. (2007). Cables links Robo-bound Abl kinase to N-cadherin-bound beta-catenin to mediate Slit-induced modulation of adhesion and transcription. Nat. Cell Biol. 9, 883–892. 10.1038/ncb1614.

104. Fogerty, F.J., Juang, J.L., Petersen, J., Clark, M.J., Hoffmann, F.M., and Mosher, D.F. (1999). Dominant effects of the bcr-abl oncogene on Drosophila morphogenesis. Oncogene 18, 219–232. 10.1038/sj.onc.1202239.

105. Dorey, K., Engen, J.R., Kretzschmar, J., Wilm, M., Neubauer, G., Schindler, T., and Superti-Furga, G. (2001). Phosphorylation and structure-based functional studies reveal a positive and a negative role for the activation loop of the c-Abl tyrosine kinase. Oncogene 20, 8075–8084. 10.1038/sj.onc.1205017.

106. Hantschel, O., and Superti-Furga, G. (2004). Regulation of the c-Abl and Bcr-Abl tyrosine kinases. Nat. Rev. Mol. Cell Biol. 5, 33–44. 10.1038/nrm1280.

107. Plattner, R., Kadlec, L., DeMali, K.A., Kazlauskas, A., and Pendergast, A.M. (1999). c-Abl is activated by growth factors and Src family kinases and has a role in the cellular response to PDGF. Genes Dev. 13, 2400–2411.

108. Stevens, T.L., Rogers, E.M., Koontz, L.M., Fox, D.T., Homem, C.C.F., Nowotarski, S.H., Artabazon, N.B., and Peifer, M. (2008). Using Bcr-Abl to examine mechanisms by which abl kinase regulates morphogenesis in Drosophila. Mol. Biol. Cell 19, 378–393. 10.1091/mbc.e07-01-0008.

109. Colicelli, J. (2010). ABL Tyrosine Kinases: Evolution of Function, Regulation, and Specificity. Sci. Signal. 3, re6. 10.1126/scisignal.3139re6.

110. Luttman, J.H., Colemon, A., Mayro, B., and Pendergast, A.M. (2021). Role of the ABL tyrosine kinases in the epithelial–mesenchymal transition and the metastatic cascade. Cell Commun. Signal. 19, 59. 10.1186/s12964-021-00739-6.

111. Zhu, G., Decker, S.J., Mayer, B.J., and Saltiel, A.R. (1993). Direct analysis of the binding of the abl Src homology 2 domain to the activated epidermal growth factor receptor. J. Biol. Chem. 268, 1775–1779. 10.1016/S0021-9258(18)53920-X.

112. Tanos, B., and Pendergast, A.M. (2006). Abl Tyrosine Kinase Regulates Endocytosis of the Epidermal Growth Factor Receptor *. J. Biol. Chem. 281, 32714–32723. 10.1074/jbc.M603126200.

113. Zak, N.B., Wides, R.J., Schejter, E.D., Raz, E., and Shilo, B.Z. (1990). Localization of the DER/flb protein in embryos: implications on the faint little ball lethal phenotype. Development 109, 865–874. 10.1242/dev.109.4.865.

114. Shilo, B.-Z. (2005). Regulating the dynamics of EGF receptor signaling in space and time. Development 132, 4017–4027. 10.1242/dev.02006.

115. Mandal, L., Banerjee, U., and Hartenstein, V. (2004). Evidence for a fruit fly hemangioblast and similarities between lymph-gland hematopoiesis in fruit fly and mammal aorta-gonadal-mesonephros mesoderm. Nat. Genet. 36, 1019–1023. 10.1038/ng1404.

116. Grigorian, M., Mandal, L., Hakimi, M., Ortiz, I., and Hartenstein, V. (2011). The convergence of Notch and MAPK signaling specifies the blood progenitor fate in the Drosophila mesoderm. Dev. Biol. 353, 105–118. 10.1016/j.ydbio.2011.02.024.

117. Dragojlovic-Munther, M., and Martinez-Agosto, J.A. (2013). Extracellular matrix-modulated Heartless signaling in Drosophila blood progenitors regulates their differentiation via a Ras/ETS/FOG pathway and target of rapamycin function. Dev. Biol. 384, 313–330. 10.1016/j.ydbio.2013.04.004.

118. Mele, S., and Johnson, T.K. (2020). Receptor Tyrosine Kinases in Development: Insights from Drosophila. Int. J. Mol. Sci. 21, 188. 10.1016/j.copbio.2018.02.003.

119. Vivekanand, P. (2018). Lessons from Drosophila Pointed, an ETS family transcription factor and key nuclear effector of the RTK signaling pathway. genesis 56, e23257. 10.1002/dvg.23257.

120. Sun, J., Ramos, A., Chapman, B., Johnnidis, J.B., Le, L., Ho, Y.-J., Klein, A., Hofmann, O., and Camargo, F.D. (2014). Clonal dynamics of native haematopoiesis. Nature 514, 322–327. 10.1038/nature13824.

121. Rodriguez-Fraticelli, A.E., Wolock, S.L., Weinreb, C.S., Panero, R., Patel, S.H., Jankovic, M., Sun, J., Calogero, R.A., Klein, A.M., and Camargo, F.D. (2018). Clonal analysis of lineage fate in native haematopoiesis. Nature 553, 212–216. 10.1038/nature25168.

122. Owusu-Ansah, E., and Banerjee, U. (2009). Reactive oxygen species prime Drosophila haematopoietic progenitors for differentiation. Nature 461, 537–541. 10.1038/nature08313.

123. Dumstrei, K., Wang, F., Shy, D., Tepass, U., and Hartenstein, V. (2002). Interaction between EGFR signaling and DE-cadherin during nervous system morphogenesis. Dev. Camb. Engl. 129, 3983–3994.

124. Rübsam, M., Mertz, A.F., Kubo, A., Marg, S., Jüngst, C., Goranci-Buzhala, G., Schauss, A.C., Horsley, V., Dufresne, E.R., Moser, M., et al. (2017). E-cadherin integrates mechanotransduction and EGFR signaling to control junctional tissue polarization and tight junction positioning. Nat. Commun. 8, 1250. 10.1038/s41467-017-01170-7.

125. Ramírez Moreno, M., and Bulgakova, N.A. (2022). The Cross-Talk Between EGFR and E-Cadherin. Front. Cell Dev. Biol. 9.

126. Bremm, A., Walch, A., Fuchs, M., Mages, J., Duyster, J., Keller, G., Hermannstädter, C., Becker, K.-F., Rauser, S., Langer, R., et al. (2008). Enhanced Activation of Epidermal Growth Factor Receptor Caused by Tumor-Derived E-Cadherin Mutations. Cancer Res. 68, 707–714. 10.1158/0008-5472.CAN-07-1588.

127. Zandi, R., Larsen, A.B., Andersen, P., Stockhausen, M.-T., and Poulsen, H.S. (2007). Mechanisms for oncogenic activation of the epidermal growth factor receptor. Cell. Signal. 19, 2013–2023. 10.1016/j.cellsig.2007.06.023.

128. Wee, P., and Wang, Z. (2017). Epidermal Growth Factor Receptor Cell Proliferation Signaling Pathways. Cancers 9, 52. 10.3390/cancers9050052.

129. Hicks, M.R., and Pyle, A.D. (2022). The emergence of the stem cell niche. Trends Cell Biol. 0. 10.1016/j.tcb.2022.07.003.

130. Sinenko, S.A., Shim, J., and Banerjee, U. (2011). Oxidative stress in the haematopoietic niche regulates the cellular immune response in Drosophila. EMBO Rep. 13, 83–89. 10.1038/embor.2011.223.

131. Konturek-Ciesla, A., and Bryder, D. (2022). Stem Cells, Hematopoiesis and Lineage Tracing: Transplantation-Centric Views and Beyond. Front. Cell Dev. Biol. 10.

132. Lee, P.-T., Zirin, J., Kanca, O., Lin, W.-W., Schulze, K.L., Li-Kroeger, D., Tao, R., Devereaux, C., Hu, Y., Chung, V., et al. (2018). A gene-specific T2A-GAL4 library for Drosophila. eLife 7. 10.7554/eLife.35574.

